# 96 Eyes: Parallel Fourier Ptychographic Microscopy for high-throughput screening

**DOI:** 10.1101/547265

**Authors:** Antony C. S. Chan, Jinho Kim, An Pan, Han Xu, Dana Nojima, Christopher Hale, Songli Wang, Changhuei Yang

## Abstract

We report the implementation of a parallel microscopy system (96 Eyes) that is capable of simultaneous imaging of all wells on a 96-well plate. The optical system consists of 96 microscopy units, where each unit is made out of a four element objective, made through a molded injection process, and a low cost CMOS camera chip. By illuminating the sample with angle varying light and applying Fourier Ptychography, we can improve the effective brightfield imaging numerical apertuure of the objectives from 0.23 to 0.3, and extend the depth of field from ±5μm to ±15μm. The use of Fourier Ptychography additionally allows us to computationally correct the objectives’ aberrations out of the rendered images, and provides us with the ability to render phase images. The 96 Eyes acquires raw data at a rate of 0.7 frame per second (all wells) and the data are processed with 4 cores of graphical processing units (GPUs; GK210, Nvidia Tesla K80, USA). The system is also capable of fluorescence imaging (excitation = 465nm, emission = 510nm) at the native resolution of the objectives. We demonstrate the capability of this system by imaging S1P_1_-eGFP-Human bone osteosarcoma epithelial (U2OS) cells.

## Introduction

The multi-well plate reader is extensively used in large format cell culture experiments. A typical well plate reader is designed to operate on either a 96-well or 384-well plate, and is usually used to perform fluorescence or absorbance measurements on the contents of the wells. These measurements can be performed quickly – readers with a throughput rate of 1 well plate per 10 seconds are commercially available^1, 2^. However, by their nature, well plate readers can only provide gross characterization of the samples.

To address this deficit, a number of imaging well plate microscopy systems have been developed over the past two decades^3–5^. These systems are typically designed to use a single microscope column to scan the entire well plate. The ability to collect microscopy images of the cell cultures provides a wealth of information that simple well plate reader cannot offer. With such a microscopy system, individual cells within a culture can be examined for their individual morphology, integrity, vitality, and their connections to neighboring cells. The range of experiments can be further expanded with the addition of fluorescence imaging functionality. For example, fluorescence imaging provides the capability to track gene expression in individual cells through the use of fluorescence biomarker methods^6^. Well plate imagers do come with a significant compromise. A single scanning microscope column has a finite data throughput rate that is set by the camera data rate and the mechanical scanning speed of the scanner itself. As a reference point, the state-of-the-art commercial well plate imager operating at an optical resolution of 1.2μm (corresponds to a typical 20× objective) can scan a complete well plate in 8min. Yet, in a per plate processing speed comparison, the per plate processing time of a well plate imager is ≈ 50 times longer than that of a non-imaging well plate reader.

The paper reports on our work at addressing this per plate throughput gap between a well plate reader and a well plate imager through the use of parallel imaging. The idea of using 96 objectives to simultaneously image all the wells on a 96 well plate may seem like an obvious and straightforward way to boost a well plate imager throughout. However, a more detailed analysis reveals several difficult engineering challenges that need to be overcome. First, ensuring that all the wells are simultaneously in focus during the imaging process is a very tall order, as each multi-well plate has its own unique warps. While the warp may be gradual, their cumulative effect across the wells can easily put a significant fraction of the wells outside the depth of field of the objectives. Second, the physical size of the objectives for parallel 96-well imaging is necessarily restricted due to their closely packed geometry in the imaging system – each objective cannot be larger than 6mm in diameter. Designing a scientific-quality objective with this size constraint can be a daunting task. Worse, each objective can be expected to have slightly different aberration characteristics due to variance in the manufacturing process. In principle, one can conceivably solve some of these problems by tightening the manufacturing tolerances and reducing residual aberrations by using more optical surfaces in the objective design. However, these engineering fixes would sharply drive up the optical system cost, especially when one considers the fact that 96 such objectives are needed in total. Third, further compounding the aberration issue, the slight difference in well curvature and well-to-lens distance for each well in a well plate introduces additional aberration variations to the imaging problem for each well. In aggregate, performing parallel well plate imaging in a conventional microscopy context is particularly challenging due to the various aberrations involved in the task.

Our approach to parallel well imaging implementation departs from the conventional microscopy system design. Here, we rely heavily on the Fourier ptychographic microscopy (FPM) method to overcome the three challenges we have just discussed.

The FPM method operates by collecting a sequence of transmission images of the sample through a microscope objective. The orientation of the illumination is varied between each image. Post image acquisition, we then computationally stitch the collected image data together in the spatial frequency domain using the Fourier ptychographic phase retrieval algorithm^7, 8^. A higher resolution image of the sample can then be recovered by inverse Fourier transform of the fused spatial frequency spectrum. The resolution of the final image is related to the sum of the objective numerical aperture and illumination numerical aperture^7^. As such, it is possible for the FPM rendered image’s resolution to be significantly improved over the native resolution of the objective^7^.

The FPM approach neatly solves the unique challenges of high-throughput imaging on a multi-well plate format described above, namely (i) defocus due to plate-to-plate variation; (ii) variation of lens aberrations; and (iii) variation of cell culture conditions. FPM offers the unique ability to characterize the aberrations of the optical system on the fly, and computationally nullify their impact on the final rendered images. We have previously shown that FPM can correct up to the 30 coefficients of the Zernike polynomial decomposition^9^. As such, we can expect the aberration challenges associated with the conventional microscopy to be significantly diminished. In addition, FPM is also effective at resolving the out-of-focus issues caused by the variation of plate geometry, as it can be considered as one of the aberrations (the fifth coefficient of the Zernike decomposition). In effect, the FPM technique can extend the effective depth of field of the imaging system beyond the limits dictated by the objective. In one of our previous projects equipped with a 2× objective (0.08 numerical aperture; native depth of field of 80μm), we showed that FPM can render in-focus images over an effective depth of field of around 300μm^7^, thereby enabling all wells to simultaneously be within focus.

In addition to solving these challenges, the FPM approach also brings the following advantages: (i) FPM’s inherent ability to improve resolution allows us to perform large field of view (FOV) imaging of cell cultures at improved resolution. (ii) Computational refocusing^7^ and aberration correction^9^ can occur post data acquisition. This is a marked benefit as users of conventional microscopy systems generally have no recourse for image correction post acquisition. (iii) The FPM image data contains phase information and we can render phase images of the sample with ease. This enables rapid, live, and label-free imaging of cell cultures.

We have previously demonstrated that the strategy of using FPM to implement parallel microscopy is feasible for 6 well plate imaging^10^. In this work, we tackle parallel imaging of 96-well plates, hence the name of our prototype – the *96 Eyes* system. Different from our previous 6 well version where we use research-grade objectives with near identical aberrations characteristics, we custom-designed our own objectives due to the high packing density required for 96 well simultaneous imaging. These objectives exhibit significant aberration variations due to surface defects from the platic injection molding process, as well as the lens alignment errors from the assembly process. As such, the completed system relies much more significantly on the robustness of the FPM technique in order to work. The 96 Eyes system also requires significant engineering work on the data transfer and processing as the data throughput rate is much higher. During operation, the system prototype acquires and transfers the raw data at 340MB/s. The system captures a batch of 64 frames of all the wells on the well plate in 90 seconds, equivalent to an imaging coverage of 3.6mm^2^ per second. Equipped with a graphical processing unit (GPU) array to accelerate FPM rendering of phase images, we can achieve the effective plate-to-image pixel rate of 2 × 10^6^ pixel/s. This system can also support fluorescence imaging at the native resolution of the objectives.

In this paper, we will first describe the design considerations that led to the current 96 Eyes system implementation. Next, we will report on the specifications of the objectives, with our experimental findings on the aberration variations between the objectives. Then, we will report on the imaging process. The data acquisition process is complicated by the variations in the well plate, i.e. warp and the menisci of the fluid within the wells. We will report on our findings and the strategies adopted to resolve the challenges. The data processing pipeline and the computational hardware and software implementations are reported in the *Supplementary Information*. Last, we report our demonstrations of our 96 Eyes system to acquire images from fixed mammalian cell lines.

## Working principle

### Design considerations

The system schematics of the 96 Eyes system is shown in Figure 1. The FPM illumination is provided by an LED matrix at the top of the system. The light transmits through the target 96 well plate. The transmission through each well is collected through an objective and projected onto a camera sensor chip. The 96 objectives and camera chips are housed on a customized 96-in-1 sensor board. The camera sensor board is interfaced with four frame grabber boards, which in turn connect to the workstation. A piezo-electric *z*-axis stage is used to hold the well plate in place and to provide *z*-axis translation as needed.

**Figure 1.**
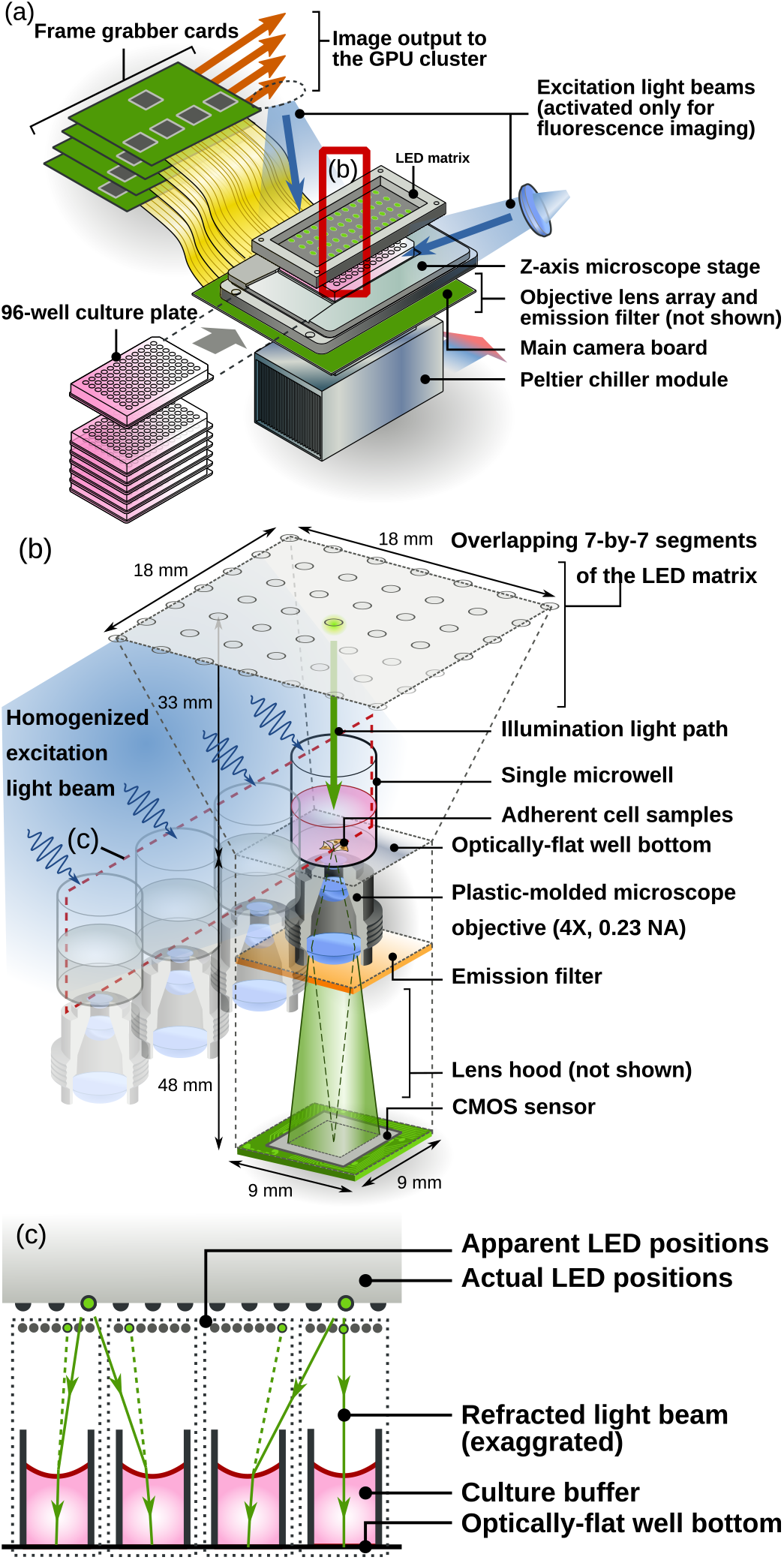
Schematic of the imaging module of the 96 Eyes system. (a) General hardware. Individual plates are loaded from the front. Refer to *Supplementary Information* for a more detailed electronic layout. (b) The imaging module consists of 96 repeating units of compact miniaturized microscopes packed in a 9mm × 9mm × 81mm space, where they all share the same light source. (c) Parallel illumination scheme revealed in the cross-section of the 96-well plate and the LED array. Refer to *Supplementary Information* for a more in-depth illustration of the parallel illumination scheme.

The 96-in-1 image sensor board incorporates 96 individual sets of CMOS sensors and microscope objectives, each set aligned to the corresponding well of the 96-well plate [Fig. 1(b)]. This arrangement results in a self-repeating units of compact microscopes in a 9mm × 9mm × 81mm space. To assemble individual CMOS sensors in such a tight layout, we choose consumer-grade CMOS sensors for their smaller footprint instead of the scientific-grade sensors. The former is also more cost-effective (up to four-fold cheaper) considering the large number (96) of sensors to be packed into the system.

To maximize the imaging FOV without sacrificing the native image resolution, the microscope objectives are custom-designed to provide a 4× magnification, a working distance of 4mm, and a numerical aperture (NA) of 0.23. The NA of 0.23 is comparable to that of a standard 10X objective (e.g. Newport M-10X). The unusual combination of a relatively high NA and low magnification is made possible in our design by the use of a much shorter tube length (34mm) by design. Notably, a finite conjugate optical configuration is chosen for its compactness and mechanical stability, resulting in a fixed object-to-image distance of 48mm. Owing to the large quantity of objectives in our system, we tolerate the manufacture imperfection of the plastic-molded lenses for its cost-effectiveness over the polished individual glass lenses or the lens array in single piece. As we will report in the lens quality characterization below, the len-to-lens variation as well as the intrinsic mis-alignment of the lens assemblies will play a role in the variation of system aberrations. We also demonstrate that the FPM method is able to retrieve and correct these aberrations.

The above described optical scheme provides an effective imaging area of 1.1mm × 0.85mm with lateral resolution of 0.61λ/NA ≈ 1.6μm, and an axial resolution of λ/NA^2^ ≈ 10μm at λ = 533nm (Table 1), supporting a native space-bandwidth product of ≈ 687 (lateral) for the fluorescence imaging light path. For phase imaging, the resolution is improved through the use of the FPM method.

**Table 1.** Design specification of the 96 Eyes system.

| Channels | Raw intensity | FPM Phase | Fluorescence |
| --- | --- | --- | --- |
| Resolution | $1.6\mu\text{m} \sim 2.0\mu\text{m}$ | $1.2\mu\text{m}^a$ | $2.0\mu\text{m}$ |
| Pixel size | $0.4\mu\text{m}$ | $0.4\mu\text{m}$ | $0.8\mu\text{m}^b$ |
| Depth-of-focus | $\pm 5\mu\text{m}$ | $\pm 15\mu\text{m}$ | $\pm 5\mu\text{m}$ |
| Field-of-view | $1.1\text{mm} \times 0.85\text{mm}$ | Same | Same |
| Acquisition time | 0.03 min | 1.5 min | $0.2\text{min}^c$ |
| Throughput | $40.0\text{mm}^2/\text{min}$ | $0.6\text{mm}^2/\text{min}$ | $4.0\text{mm}^2/\text{min}$ |

<sup>a</sup>The lateral resolution is influenced by the deviation from ideal object-to-sensor distance. Refer to Fig. 2(b) for detail.
<sup>b</sup>Pixel binning is applied to reduce noise.
<sup>c</sup>Assuming digital averaging of 10 frames and z-scanning only one layer.

FPM image acquisition involves illuminating the object with a sequence of spatially distributed light sources. In order to make sure all specimens on the 96-well plates receive an identical set of illumination conditions, we incorporate a large area light-emitting diode (LED) array with the LED-to-LED separation of 3mm, exactly one quarter of the well-to-well separation of the 96-well plate. The vertical clearance of the LED array is chosen to be at 33mm from the sample plane, so as to ensure sufficient redundancy of the collected ptychographic data^11^. For ptychographic imaging, each repeating unit of the mini-microscope effectively utilizes an 18mm by 18mm segment of the LED array. In the tight layout of the mini-microscopes (9mm by 9mm), exactly three quarters (75%) of the area of a LED array segment are reused by the adjacent mini-microscopes. As a result, the total number of LED elements required for the 96 Eyes system (= 1120) is reduced by 75% (from 96 × 7^2^ = 4704). Such a high density layout also enables us to illuminate all wells on the 96-well plate by turning on multiple LEDs. The working wavelength is chosen to be ≈ 530nm, matching the passband of the fluorescence emission filter [Fig. S8]. The implementation scheme will be described in detail below.

### Parallel illumination scheme

Our image acquisition approach harnesses the fact that the illumination system is shared among multiple imaging sensors, each of which captures individual specimens independently [Figs. 1(b–c)]. In the case of an array level Fourier ptychographic imaging system, an individual specimen corresponds to a discrete well of the multi-well plate. Here, we utilize the high density of the 96-well plate, where a single LED can illuminate up to nine wells at the same time. We further exploit our design specification (NA_target_ ≤ 2NA_lens_) to get rid of all darkfield image recording on the fly. This allows us to turn on multiple LEDs to fully illuminate the entire 96-well plate, while making sure that each well is only illuminated by a single light source at any time. This parallel approach allows us to minimize camera idling time [Fig. 1(c) and Supp. Fig. S2]. Therefore, image acquisition and data transfer can be performed in a massively parallel manner during the imaging step.

To avoid superposition of any two light sources on the same well, we generate a rectangular grid of illumination pattern on the LED array with a source-to-source separation of *m* LEDs. The value of *m* is chosen to be large enough so that for any wells at any time instance, only one LED is located within the light acceptance cone of the corresponding microscope objective determined by the numerical aperture [Supp. Fig. S2]. Refer to Supp. Fig. S2 for more detailed illustration of the parallel illumination scheme.

To further suppress stray light from an adjacent lens impinging on the image sensor, a wafer-shaped lens hood array surrounds and shields the light path between the objectives and the CMOS sensors [Fig. 1(b)]. This light shielding can be especially important in fluorescence imaging where the intensity of the emitted light is much lower than that of the excitation light.

Similar to the parallel image acquisition scheme, a multiple image restoration algorithm is performed in parallel, as all sensors receive an identical set of illumination patterns [Figs. 1(b–c)].

## Results

### Characterization of aberrations of plastic-molded microscope objectives

The ultimate resolution performance of the 96 Eyes system hinges on the manufacture tolerance of the custom-designed objectives. Variations of the lens element geometry, as well as relative mis-alignment of elements within the objective barrel, can introduce varying amounts of aberrations among different samples, and can in turn introduce statistical errors in the experiments carried out in the 96-well plate format. This is especially important in fluorescence imaging mode in native resolution determined by the corresponding objectives.

Our choice of finite-conjugate objectives also requires a very specific object-to-sensor distance. To evaluate the effect of distance deviation on the lens aberrations, we used manufacturer provided simulation data of the spot diagram when the distance shifts by (i) ±2mm and (ii) ±5mm at a wavelength of 530nm [Fig. 2(b)]. The object-to-objective-barrel distance is allowed to adjust to ensure maximum resolution. The data indicates that the spot size can stay within the disk of 2.5μm diameter when the sample-to-sensor distance stays within ±2mm of the ideal distance at 48mm. This implies the bottom side of the 96-well plate has to be almost perfectly parallel to the sensor board, with a relative tilt tolerance as small as arctan[±2mm/(9mm × 11)] ≈ ±1°. We tackle the low tolerance constraint by precisely aligning the mounting bracket of the microwell plate, as well as choosing a more rigid material for the plate itself.

**Figure 2.**
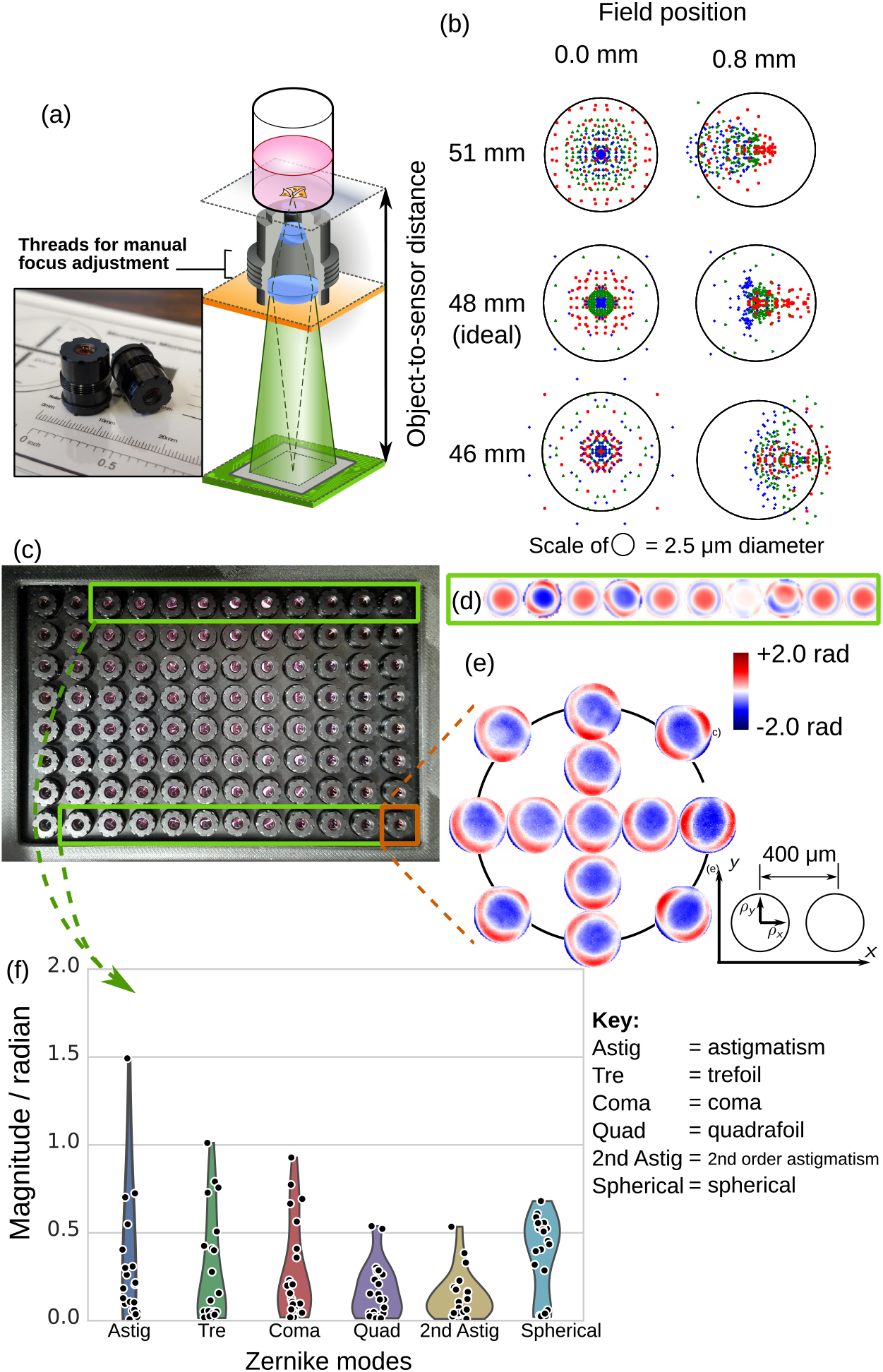
Characterization of lens aberration in 96 Eyes system. (a) Illustration of the object-to-sensor distance for finite conjugate optics. The external threads of 350μm pitch allows manual adjustment of lens-to-object distance. The inset shows the actual scale of the plastic-molded microscope objectives. (b) Spot diagram simulation of the lens at on-axis (0μm) and off-axis position (800μm), when the object-to-sensor distance deviates from design value (i.e. at 48mm). In the simulation, the objective lens was allowed to move to optimize the focus. (c) Top view of the custom-designed microscope objectives assembled in array format. (d) Aberration of ten (10) microscopes are recovered by FPM-EPRY algorithm. (e) “Local” pupil function recovered from FPM-EPRY algorithm, showing a slight spatial dependency of the aberration. (f) Distribution of the rotation-invariant Zernike aberration modes of 21 microscopes, measured from the center of FOV.

Next, we sampled 21 objectives from our 96 Eyes system [Fig. 2(c)], and measured the aberrations of multiple objectives with the optimal object-to-sensor distance. Refer to Materials and methods for detailed system configuration and sample preparation. We adopted the FPM for optical profiling for its effectiveness in measuring lens aberrations *in situ*, without resorting to additional optical instruments for wavefront sensing^12^.

The local lens aberrations were measured from the center of the image FOV and were fitted with the first 15 Zernike modes [Fig. 2(a)]. We found that the pupil function exhibits a variation due to manufacturing variation of plastic-molded lenses, with a standard deviation of 0.88 radian on astigmatism, and 0.25 radian on spherical aberration. Figure 2(d) exhibits the pseudo-color phase map, with piston, tilt, and defocus mode removed for clarity. There is also a significant off-axis aberration, with a root-mean-square error of 0.54 radian compared to that at the center of FOV. Again this is captured by the local Zernike fit of the field position of 0.4μm and 0.8μm respectively [Fig. 2(f)].

To ensure the flatness of our sample, we also probe the sample surface with our standalone FPM microscope retrofitted from a scientific-grade widefield microscope [Supp. Fig. S6]. The surface variation of the sample is found to be below 0.35 radian, below the detection limit of our scientific-grade FPM microscope. To ensure the consistency of our measurements, we also prepared a separate microbead sample and repeated our measurements.

### Characterization of well depth variation

Although the FPM is capable of providing an extended depth-of-focus in the final rendered images, it is nevertheless desirable for the well plates themselves to have a consistent physical depth profile. Among various commercially available choices, we picked the Grenier UV-Star plate because its well bottoms are made of a more rigid material (cyclic olefin copolymer, COC) then those found in geneic culture plates, which are made of either polystyrene or polypropylene.

We characterized the depth variations by analyzing 16 plates (UV-Star plates) using an Opera Phenix high content imaging system equipped with laser-based autofocus. For comparison, we also evaluated 16 generic culture plates with polystyrene bottoms (Cell Star plates). As shown in Figs. 3(a–b), both plate types exhibit a general curvature of the well bottom with a depression of up to 500μm. A general tilting of the well bottom is also observed in the depth profile; the wells appear to be more depressed (consistently across all plates) at the bottom left corner. This may imply a slight tilting of the sample plane relative to the flange of the 96-well plate where the former sits on. Such *per well* average tilting and curvature can be compensated with a one-off manual focusing step for individual microscope objectives.

**Figure 3.**
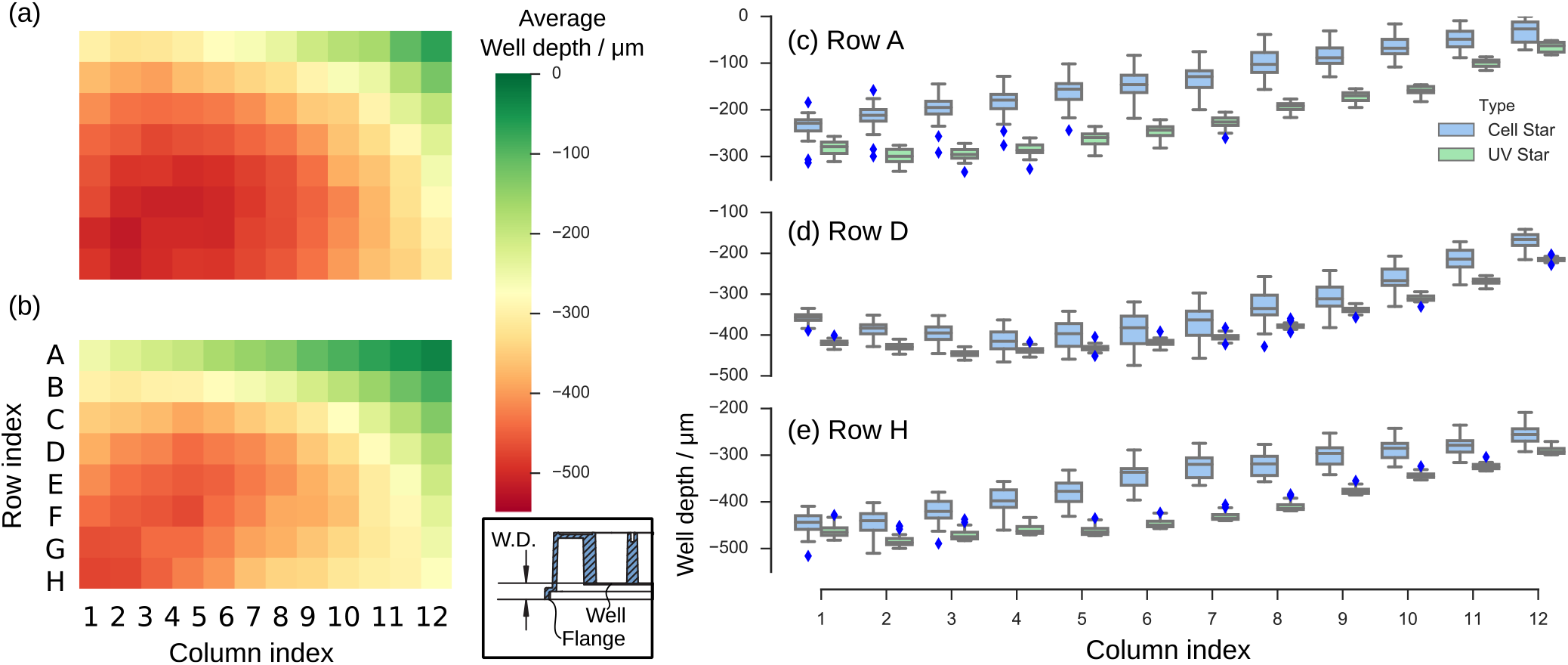
Depth measurement of two different models of the Greiner 96-well cell culture plates. Average well depth of 16 units of (a) Cell Star, and (b) UV Star. Box plot of the well depth distribution along (c) Row A; (d) Row D; and (e) Row H. Inset: The well depth (W. D.) is defined as the vertical distance between the flange of the 96-well plate and the top side of the well bottom. For the sake of clarity, the depth measurements are subtracted from that of the shallowest well of all plates.

The well depth distributions are also analyzed along the top row, middle row and the bottom row [Figs. 3(c–e)]. The COC-bottom plates shows a much tighter depth tolerance (range ≤ 66μm, standard deviation ≤ 17μm) across all 16 similar plates, compared to that of the generic polystyrene-bottom plate (range ≤ 170μm, standard deviation ≤ 45μm).

### Characterization of extended depth-of-focus enabled by FPM

As revealed in the above study, natural geometrical warping of the cell culture plates is significant enough that a conventional plate imager would have to compensate by mechanical depth scanning and refocusing. Such a significant plate-to-plate variation of up to ±33μm is more than 3 times the native depth resolution (λ/NA^2^ ≈ 10μm at λ = 533nm) permitted by the microscope objectives [Supp. Fig. 3]. To this end, we characterize the robustness of FPM in restoring the naturally out-of-focus images. Here, we intentionally and precisely shifted the sample plane of a Siemen Star phase target followed by FPM acquisition [Fig.4(a)]. The captured data undergo FPM reconstruction with computational refocusing^12^. Specifically, the wavefront error associated with the focal shift was blindly estimated by minimizing the FPM reconstruction residual [Eq. (S15) in *Supplementary Information*] of the refocused image with a linear search algorithm. After computational refocusing, the phase quality of the FPM output is evaluated by measuring the lateral resolution. Refer to Materials and methods for detailed experimental condition and sample preparation.

**Figure 4.**
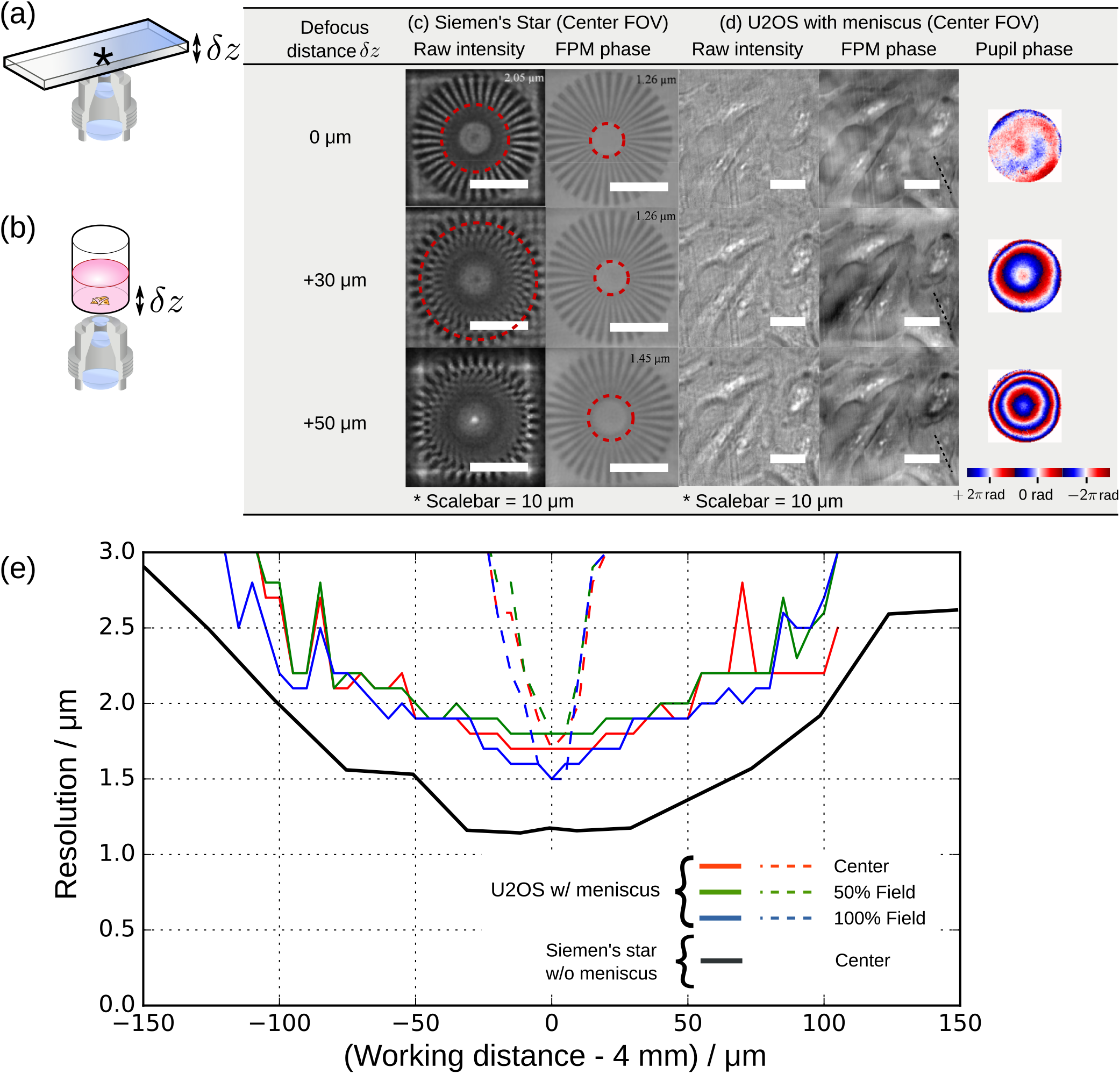
FPM enables extended depth-of-focus microscopy of up to ±50μm, reducing the time spent on mechanical focusing. To characterize the range of the depth-of-focus permitted by our 96 Eyes system, we perform *z*-stack mechanical scanning to introduce out-of-focus aberration intentionally for (a) the phase-only Siemen’s star target, and (b) the U2OS cell line with liquid meniscus, (c) Plot of the lateral resolution of the digital refocused image. over a range of [-150μm, +150μm]. The dotted line in the same plot illustrates the resolution of the phase image if the computational refocusing step is omitted. (d) The table shows the side-by-side comparison of the raw image and the corrected phase image, as well as the recovered phase component of the pupil.

The restored Siemen’s star phase images are compared against the raw intensity image captured from illumination of a single LED [Fig. 4(c)]. At ideal working distance from the microscope objectives (i.e. *δz* = 0μm), the lateral resolution of the FPM restored image is ≤ 1.26μm, below the detection limit of the Siemen’s star target. In contrast, the native lateral resolution of the same pattern is 2.05μm, in agreement with the theoretical resolution of the intensity image. When the object is axially shifted by *δz* = +30μm, we observe a severe distortion of all spatial features below 3.5μm. But with the computational refocusing method, the restored phase image is almost perfectly restored at resolution ≤ 1.26μm. Again, this is below the detection limit of the resolution target. The phase image resolution only degrades marginally at 1.45μm at an axial shift *δz* = +50μm. The experiments were repeated over the range of −150μm ≤ *δz* ≤ +150μm at 25μm step size, which results in the lateral resolution versus defocus distance plot [black trace in Fig. 4(e)]. The restored lateral resolution of the phase image stays at around 1.26μm for the defocus range of [−30μm, +30μm], then ramps up to around 1.5μm at *δz* = ±50μm, and around 2.0μm at *δz* = ±100μm.

We performed a similar experiment with an adherent cell line seeded and fixed in a 96-well plate [Fig. 4(c)]. Refer to Materials and methods for the cell preparation protocol. The raw intensity image and the corresponding restored phase image at corresponding depths (*δz* = 0μm, +30μm, +50μm) are also tabulated in Fig. 4(c). Here, an axial shift of |*δz*| ≥ 30μm is sufficient enough to severely degrade the raw intensity image. We show that the unknown defocus distance can be blindly estimated by minimizing the FPM reconstruction residual with the linear search algorithm. This additional step is particularly important for accurate phase image recovery, as the embedded pupil recovery algorithm alone is less likely to converge to a global minimum^13^ when the defocus distance is beyond the depth-of-focus (10μm), at which the phase difference in the wavefront error can exceed 2*π*. With the two-stage computational refocusing method, the cell images are successfully brought back to focus. The recovered aberration maps, also presented in Fig. 4(c), exhibits a quadratic phase profile which indicates a significant contribution of defocus aberration then other higher order aberration modes. We also applied computational refocusing at off-axis positions [Supp. Fig. S7], revealing consistent phase image quality as well as aberration map.

To characterize the tradeoff between the phase image quality of adherent cell samples and the defocus distance, the experiment was repeated over a range of −120μm ≤ *δz* ≤ +120μm at a step size of 5μm. The lateral resolution is estimated by measuring the full-width-at-half-maximum (FWHM) of the line profile across selected granules within the cell [black dotted line in Fig. 4(d)]. The restored phase image resolution stays below 2.0μm when the defocus distance |*δz*| is below 50μm. In contrast, the phase image resolution without computational refocusing degrades significantly (≫ 2.0μm) as soon as the cell sample is shifted beyond 15μm [dotted line in Fig. 4(e)].

### Calibration of illumination angles through thick liquid medium

Embedded pupil function recovery algorithm^9^ enables aberration correction of the image *in situ*. However, it requires accurate estimation of the LED positions above the samples, as well as the angle of incidence on the sample through the liquid medium. Conventional cell culture protocols require that the samples, either living cells or fixed cells, be hydrated with culture buffer of at least 100 μL (for samples in a 96-well plate), which translates to around 3mm thickness of liquid medium. This thick liquid medium act as a fish-eye lens that alters the beam of illuminations from the LEDs to the sample, which introduces errors in FPM phase recovery. To compensate this effect, we introduced the ray tracing algorithm based on Snell’s law. The detail ray tracing framework is included in the *Supplementary Information*. Notably, such a precise calibration step was not crucial in our previous designs of FPM imagers on standard microscope slides^7^, or on a 6-well plate^10^, where the medium is flat and the is thinner than 1mm.

The challenge to estimate illumination angles through liquid medium is further compounded by the fact that the liquid medium surface forms a curved meniscus surface in the 96-well plate – it acts as a concave lens that brings the image of the LEDs much closer to the sample. This meniscus brings a challenge of ptychographic illumination, known as the parallax effect, where the position of the light source appears to differ when viewed from different lateral positions on the object plane [Fig. 1(c) and Supp. Fig. S2]. Our observations with the raw data reveals that the angle of incidence can drift by 5° from the center of FOV to the edge (i.e. a 0.5mm lateral displacement), that is 6 times of the shift without liquid meniscus. Again, we compensated for this effect with our ray-tracing algorithm as described in *Supplementary Information*.

### 96-well plate cell culture imaging results

With the 96 Eyes prototype ready, we next imaged cell cultures grown in 96 well plates. In this demonstration, we used the human bone osteosarcoma epithelial cell line (U2OS) modified to express green fluorescent protein (GFP)-tagged S1P1, or sphingosine-1-phosphate-receptor 1, which translocates from the plasma membrane to the nucleus in the presence of specific ligands. The cells were seeded, incubated overnight, then imaged after confirming cell attachment. The GFP expression is readily excitable with an external excitation source at 465nm. Refer to Materials and methods for detailed optical configurations and the cell preparation protocol.

We note that our choice of consumer-grade CMOS imaging sensors results in a relatively low image dynamic range. To enhance the signal-to-noise ratio, we digitally average multiple video frames of the same field-of-view to denoise the image. The band-like static pattern is also removed with a similar technique [Supp. Fig. S5]. Refer to *Supplementary Information* for the theoretical framework of two-stage digital averaging technique. With this digital frame averaging technique, we are able to capture fluorescent signals from the whole plate within 10s (Table 1).

Figure 5 shows the resulting images as collected with the 96 Eyes. Figure 5(a) shows the low resolution image captured from all 96 microscopes trained on the corresponding wells on the 96-well plate. One of the wells is post-processed with our GPU-accelerated FPM algorithm (refer to) to restore the phase image, in addition to the de-noised fluorescence image selected from the *z*-stack scans [Fig. 5(b)]. To highlight the strength of FPM for phase contrast imaging of unlabelled cell samples, we picked two regions of 110μm by 110μm for closer inspection. Compared to the low resolution, out of focus raw images [Fig. 5(c)], the restored phase images [Fig. 5(e)] present the aberration-free structures of cellular morphology. For data completeness, the FPM algorithm also outputs the intensity component of the restored object [Fig. 5(d)], yet the cells are almost invisible under our working wavelength at 530nm. The de-noised fluorescence images are shown in Fig. 5(f). The morphology of the cell bodies match perfectly with the corresponding recovered phase image [Fig. 5(e)]. The composite image is also generated by fusing the phase image and the fluorescence image [Fig. 5(g)]. Notably, the two image channels align well without the need of any computational pixel registration software.

**Figure 5.**
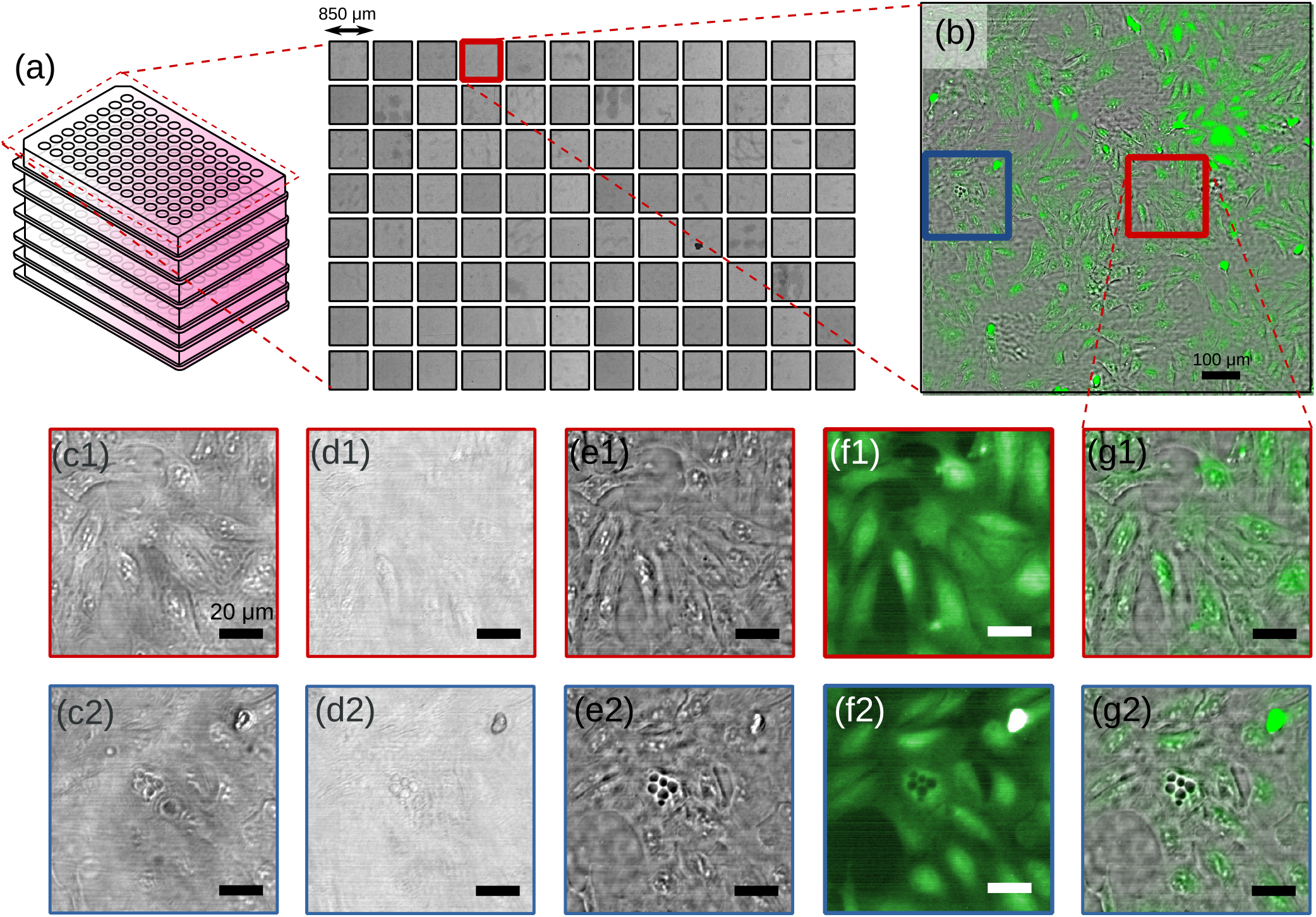
High-throughput multimodal imaging enables cell screening at a massive scale. (a) Montage of all 96 brightfield images captured by the 96 Eyes imager; (b) Magnified view of well A8 on the 96-well plate; (c) Low-resolution, out-of-focus raw image captured by the consumer-grade CMOS sensor; (d) Restored intensity of the object; (e) Restored phase; (f) Fluorescence response from eGFP; (g) Composite image of phase channel in gray color and eGFP channel in green color. The cells are S1P_1_-eGFP-Human bone osteosarcoma epithelial cells^14^ (U2OS; ATCC number HTB-96), seeded and fixed in the 96-well culture plate. Refer to Materials and methods for the detailed cell preparation protocol.

Unlike the phase image channel where the depth-of-focus (DOF) can be extended computationally with digital refocusing, the fluorescence image is limited by the native DOF imposed by the microscope objectives. Here, the plates are scanned along *z*-axis with a step size of 25μm (around twice the native depth resolution (= 10μm) permitted by the custom-designed microscope objective), and a range of 100μm (around the 90% confidence well-depth range of the UV-Star plate).

## Discussions

We characterized the lens aberrations from identically-designed microscope objectives, which reveals a variation of 0.88 radian on astigmatism and 0.25 radian on spherical aberration. Such a tight lens aberration tolerance is required for consistent measurement for multiple plates, which ultimately improves the statistical strength of high-throughput biomedical studies. This is particularly important in fluorescence imaging because the image quality is directly tied to the native image resolution dictated by the microscope objectives. Although we previously demonstrated that the fluorescence image can be deblurred with the recovered aberration from the phase image channel^15^, the method is not used in this prototyping project as the image quality is sufficient in the 96 Eyes system. The tight tolerance also improves the data analysis of phase images recovered by FPM. Even though they can be aberration-corrected computationally, a strong manufacture variation of the objectives can cast doubts on the data consistency of the images captured among different lens. For instance, we have shown in the extended depth-of-focus study that the lateral resolution can be degraded by a fraction even with accurate estimation of the aberration wavefront. Thus, it cannot be ruled out that strong values in higher order modes of aberrations can further degrade the phase images. Here, a tight manufacturer tolerance of the microscope objectives helps eliminate such doubts, and in turn improves the statistical strength for high-throughput studies with the 96 Eyes system.

The FPM-based aberration correction poses a unique strength in determining the *overall aberration* of individual objectives on the fly. For instance, it uses the exact same illumination condition and optical configuration as the phase image acquisition of cell samples. No alternation of the light path is required. More importantly, embedded aberration recovery with FPM preserves the direction of the aberration components (e.g. slopes of the tilted wavefront, major axes of the astigmatisms) of the microscope objective at the calibrated position in the 96 Eyes system. Such directional information is essential in correcting aberration of the phase images, which can be lost by merely rotating the lens barrel – a necessary step in assembly and lens-to-sensor distance calibration. Notably, the best alternative is to move the objectives from the 96 Eyes instrument to the specialized optical profiler for aberration analysis. The measured lens aberration map may have served the purpose of lens tolerance study, but has limited value in the aberration correction of phase images. To this end, we are able to compensate for the discrepancies in wavefront error to correct the phase image with our FPM technique for any positions in the image FOV.

The extended depth-of-focus study demonstrates that the variation of object-to-lens distance can be corrected with the aberration correction method within the FPM phase retrieval algorithm, which corrects the out-of-focus aberration associated with the focus offsets. As a result, we can tolerate an extended depth variation of ±50μm in the 96-well plate. Through the well-depth characterization study, we picked the UV-Star plate which fits our system tolerance, having a tighter depth range than other generic culture plates, with a standard deviation of 17μm. In other words, an extended depth-of-focus of 100μm full range can cover around 90% of all UV-Star plates. It is also shown that the effect of within-well warping is negligible compared to lens aberrations and defocus.

The large well depth variation is understandable due to the fact that 96-well culture plates were originally designed for photo-chemical analysis^16, 17^, in which optical precision is not a requirement. In spite of a number of recent attempts to re-design the multi-well plate to address the above issues^18, 19^, the 96-well plate format remains popular among researchers and thus remains the gold standard in high-throughput screening applications^17^.

Notably, the plate-to-plate depth variation concern is unique to our 96 Eyes system, as the lens positions (i.e. lens-to-sensor distance) are locked during image acquisition of multiple plates. This is in contrast to conventional plate scanning microscopes, where either the objective or the plate can be can be mechanically actuated to focus on individual wells, albeit with a significant tradeoff in speed. In our current implementation of the 96 Eyes system, we simply pick the best plate type and let the FPM phase retrieval algorithm correct the aberration associated with the plate warping. In our study, we have not looked into the connection of well depth variation to various aberration modes in the imaging light path. The choice of finite conjugate objectives dictates a fixed sample-to-image-sensor distance; any deviation from it can result in higher order aberrations other than the defocus mode. Such a study may provide a more thorough understanding of the performance of the custom designed microscope objectives. Nevertheless, this knowledge does not improve or degrade the phase image restoration, as FPM algorithm is not sensitive to the source of aberrations, whether they stem from lens manufacturing errors, lens position errors, well-depth errors, or a combination of all of the above.

It is also worth noting that on-the-fly focus adjustment of the individual microscope objectives is possible with either the *voice coil motor* (VCM)^20^, the micro-electrical-mechanical system (MEMS)^21, 22^ actuator, or the variablefocus liquid lenses^23, 24^, eliminating the plate-to-plate depth variation early at the acquisition step. So far, we have not found any commercially available solutions that can fit the 9mm × 9mm format. However, in the foreseeable future, we anticipate that micro-actuation technologies will enable 96 Eyes to accept a wider range of 96-well plates with more relaxed plate warping constraint.

As a multi-modal imaging system, 96 Eyes can image fluorescence images in visible wavelength. Because both FPM and fluorescence channels share the same imaging light path, the two images are automatically co-registered and coaligned. We mitigated the image blur in fluorescence channel by capturing a z-stack of fluorescence images over a range of ±30μm, around 5 times the native depth resolution permitted by the microscope objectives. The sharpest image was then selected from the *z*-stack.

It is noted that such a *z*-position can also assist in determining the out-of-focus distance in FPM channel, thus further improving the speed of phase retrieval and aberration correction with FPM algorithm.

The power-line electronic noise can be further suppressed with a larger value of the filtering capacitor. Our current implementation places the capacitor on the same side as the CMOS sensor, limiting the capacitance value possible for noise suppression. We are working on our next design iteration to place a larger value capacitor on the other side, with custom-designed heat sinks.

This has been demonstrated with CMOS sensors chilled to below room temperature^25^ to minimize dark current noise in the image. Our current design of the Peltier chiller is capable of chilling the consumer-grade CMOS sensors, further improving the fluorescence sensitivity. However, the modular design of our electronics, an oversight for low-cost CMOS repair, introduces poor thermal conductivity between the CMOS sensors and the bottom surface of the main PCB, and thus limits the cooling capability of the chiller. As a result, CMOS sensors run at ≈ 10 °C above the ambient temperature, and as such can contribute to a significant amount of thermal noise to the pixel readout process. Going forward, we believe that thermal conduction can be improved by soldering the sensors directly onto the PCB surface.

## Conclusions

We introduced the 96-in-1 parallel imaging system (96 Eyes) for high throughout cell screening. By using 96 repeating units of low-cost image sensors and custom designed objectives to simultaneously image all the wells on a 96-well plate, the overall imaging throughput can be improved significantly. Here we addressed and characterized the concern of lens-to-lens variations and deviations from the ideal sample-to-lens distance; both issues can result in poor image quality. We also characterized the culture plate warping by comparing two plate types, and observed its effect on the out-of-focus aberrations in the imaging system. The effect of liquid meniscus on the image quality is much stronger than anticipated, introducing additional image aberration. We heavily rely on the FPM method to overcome the above challenges, utilizing the embedded pupil recovery algorithm with adaptive step size, as well as the meniscus-compensating ray tracing method. Equipped with the custom designed 96-in-1 CMOS array board, we successfully captured ptychographic intensity images of a 96 well plate within 90 seconds, and fluorescence images within 30 seconds. The ptychographic images are processed with GPU acceleration, feeding endusers with high resolution (1.2μm), aberration-free phase contrast images of 1.1mm by 0.85mm per condition at the extended depth-of-focus of ±15μm.

We also note the potential of a better imaging robustness through hardware improvement. For instance, 96 Eyes system can be upgraded with variable-focus micro-actuators or liquid lenses to control the imaging focus independently, so that the system can accept a wider range of 96-well plates with a more relaxed plate warping constraint. In particular, meniscus-free culture plates^26, 27^ can be adopted to the 96 Eyes system for a more predictable illumination angles, thus reduces the errors in FPM phase restoration. On the other hand, the optical sensitivity to fluorescence can be increased with a better powerline noise rejection and thermal management of the 96-in-1 CMOS sensor array. A second fluorescence channel at longer wavelengths (i.e. between 560nm and 700nm) can also be included in the system with a multi-bandpass emission filter and a second excitation light source. However, since the plastic-molded microscope objectives were designed at a narrow range of wavelengths (i.e. around 530nm), further studies are required to evaluate the aberrations at longer wavelengths.

We anticipate that the 96 Eyes system will help accelerate biomedical and pharmaceutical research that utilizes high-throughput cell imaging and screening format.

## Materials and methods

### Fluorescence imaging

A pair of liquid-guided excitation sources projects the homogenized light beam (Nichia NUBM07E filtered by Semrock FF01-466/5-25, bandwidth 465nm ± 2.5nm, power 4000mW) from both sides of the culture plate. Due to the extreme angle of illumination, the direct transmitted light is blocked by the internal aperture of the microscope objectives. The residual scattered light is further attenuated by the emission filter (Chroma 59022m, center wavelength = 535nm, bandwidth = 50nm). Refer to Supp. Fig. S8 for the combined plots of the excitation wavelength, passband of the emission filter, and the quantum efficiency curve of the eGFP. To compensate focal shift, the plates are scanned with the *z*-axis piezo flexure scanning stage capable of 2.5μm repeatability and a range of 300μm (Dynamic Structures & Materials Model ZSA-300, USA). The plate is scanned with a step size of 25μm (around twice the native DOF), and a range of 100μm (around the 90% confidence well-depth range of the UV-Star plate). For a given *z* position, digital frame averaging is applied to the time sequence of captured image for all step *z* positions.

### GPU accelerated parallel FPM reconstruction

The GPU-accelerated ptychographic phase retrieval algorithm is implemented with the cuFFT and Thrust interface^28^. Illumination angle estimation is implemented with the Armadillo matrix library^29^. Acquisition and saving implemented with the MPI-ready parallel HDF5 interface (High-Five) developed by the Blue Brain Project team at EPFL^30^.

Acceleration of the FPM phase image reconstruction is made possible by utilizing the fact that all 96 cameras receives identical sequence of illumination conditions. The entire imaging area of a given well on a 96-well plate is partitioned into an 8 by 10 grid of slightly overlapping image segments of 110μm × 110μm dimensions. The segments of all 96-wells are grouped by their corresponding lateral positions, and then sent to the GPUs (GK210 processor ×4, Nvidia Tesla K80, USA) in batches for FPM reconstruction. Further acceleration is accomplished by pipelining the read-render-write process to eliminate the idling time of the GPU processors. The intermediate FPM intensity and phase images, expressed in complex values, are stitched with feather blending^31^ to recover the whole FOV of the images. The image stitching and rendering process is accelerated on a pair of 10-core CPUs and are archived on the 15 terabyte hard drive for further analysis. Refer to *Supplementary Information* for a more detailed description of the acceleration method.

### Fabrication of Siemen star phase targets

The Siemens star resolution phase target, recommended by Horstmeyer et al^32^, is fabricated on a gold-coated glass. Using focused ion beam (FIB), the Siemens star pattern is first etched onto the gold surface. Then, a 50μm × 50μm area encompassing the whole pattern is further etched with the same exposure time until the glass substrate is exposed within the entire area.

### Cell culture conditions

S1P_1_-eGFP-Human bone osteosarcoma epithelial cells^14^ (U2OS; ATCC number HTB-96) were maintained in Dulbecco’s Modified Eagle’s medium (Life Technologies) supplemented with 10% fetal bovine serum (FBS), 2mM L-Glutamine, and 1% Penicillin Streptomycin (Life Technologies). Cells are seeded at a density of 8,000 cells per well in the 96-well culture plate (Grenier UV-Star or Cell Star) and incubated overnight at 37°C and 5% carbon dioxide. For the UV-Star plate, the well bottom was first treated with Synthe-max to allow the cells to attach. The cells are then fixed and cross-linked with formaldehyde, followed by rinsing with Dulbecco’s Phosphate-Buffered Saline solution (DPBS). The fixed cells in each well are then hydrated with 200 μL DPBS solution before imaging.

### Microsphere sample preparation

Polystyrene microsphere of 2μm diameter [Polyscience Inc., USA, Cat. 19814-15] is suspended in DPBS and diluted to 53,000unit/μl. Then, 10 μL is sampled from the suspension and deposited onto a single well of 96-well plate, followed by adding a cover glass of 5mm diameter. The 96-well plate is then loaded into the 96 Eyes instrument for ptychographic imaging.

### Microwell plate well flatness

To measure the overall flatness of COC microplates (Grenier UV-Star Cat. No. 655801) compared to the polystyrene-based microplates (Grenier Cell Star Cat. No. 655160), the Opera Phenix Absorbance Microplate Reader was used to measure the distance from the flange to the well bottom. Sixteen (16) plates from two models of the 96-well plate, are seeded with cells (described above) before the measurement. The difference between the measured distance and the minimum distance of all measured plates are used to generate an overall well depth profile.

An FPM microscope, retrofitted from a standard widefield microscope (Olympus Model XL-41) equipped with a 20X objective (Olympus Plan N 20X; 0.4 NA)^7^, was used to measure the flatness within each well. The well was first seeded with polystyrene microspheres (described above) before ptychographic imaging. Five (5) locations within the selected 24 wells of a 96-well plate are measured by the FPM technique with local aberration recovery, which in turn was used to compute the defocus distance from the focal plane. The difference between the maximum and minimum value was calculated to determine the variance in flatness within each well. The variance of each well was than averaged to determine overall well flatness.

## Author contributions

Conceived and designed the hardware/software/experiments: A.C.S.C., J.K., A. P. Contributed reagents/materials/analysis tools: H.X., D.N., C.H. Wrote the paper: A.C.S.C., A.P. Supervised the project: S.W., C.Y.

## Acknowledgments

We would like to thank Daniel Martin for fabricating the Siemens star phase target, and to thank Albert Chung for the suggestion of an adaptive step size for more accurate pupil recovery. We also thank Kevin Wong and Richard Graham from ClearBridge Biophotonics for the technical advices on coding the image stitching algorithm and background interference removal.

