## Supplementary Information for "96 Eyes: Parallel Fourier Ptychographic Microscopy for high-throughput screening"

### ABSTRACT

This document provides supplementary information to the "96 Eyes: Parallel Fourier Ptychographic Microscopy for high-throughput screening," [year], pp. [page].

### Supplementary Information

#### Parallel FPM acquisition and reconstruction

A parallel image acquisition technique is proposed here. Four (4) frame grabbers are simultaneously controlled by individual processes in the workstation, each is run in individual central processor core. One of the process supervises the illumination system to implement step, and then sends out trigger signal to all other processes to perform image acquisition and storage. As shown in Figs. S1(b–c), the ratio of the number of image sensors to the number of running processes is equal to 24, that corresponds to four set of 24-to-1 multiplexers for a total of the 96 image sensors. With respect to target applications, such ratio can be varied to optimize the overall data throughput within the allowable bandwidth of the interface.

Another data throughput challenge preciously not addressed in previous studies (e.g. EmSight[1]) is the requirement of segmenting the image data into tiles on the fly. If the images are first saved and segmented later, both the imaging system and the graphical processor(s) will be idling, thus it limits the overall image restoration throughput. Our study shows that our system can finish writing the raw image data within 2 minutes, yet it takes around 20 minutes to reorganize (i.e. read, segment, and write) the raw data from/to the hard drive. This challenge can be addressed with a in-memory parallel data storage strategy accessible by all running processes, which houses a four-dimensional image data "hypercube" with a dimensions of "number of illumination angles" times "number of image sensors" times "image height" times "image width". The hypercube is pre-segmented into chunks of dimensions (in our case, it is $1 \times 96 \times 256 \times 256$). For each unique illumination pattern, the incoming image data of all image sensors are simultaneously sorted, indexed, and segmented online by the file system. The individual chunks of the hypercube are then written to the hard drive in a linear layout, which facilitates the image segment loading and restoration method in the next step. In short, by sorting and segmenting the incoming image data on the fly, it helps saving the precious data bandwidth.

Enabled by the data alignment of the image chunks and the identical illumination pattern across all image sensors, multiple image segments can be restored by the graphical processor in a massively parallel manner. The corresponding image segments for all image sensors (i.e. at identical locations in the image FOV) can be processed simultaneously as they possess an identical set of illumination conditions. This substantially reduces the GPU idling time because a chunk in the data "hypercube" only requires one set of function calls and data read/write instead of 96 (for 96 image sensors), reducing the processing overhead.

If only a single 96-well plate is imaged and analyzed, the back-to-back data acquisition (one layer of phase image plus 10 z-layers of fluorescence image) and processing pipeline requires $(90+30+120) = 240\,\mathrm{s} \approx 4\,\mathrm{min}$ to complete. However, if multiple plates are involved in one batch of study, the acquisition stages and the reconstruction stages can be performed simultaneously (Fig. S1), reducing the overall imaging time to around 120 second per plate.

#### LED position calibration

Fourier ptychographic algorithm requires accurate illumination angles from different LEDs in order to register the raw images in the Fourier domain. Because of the presence of liquid meniscus in the 96-well culture plate, the LEDs appears to be much closer to the object than they are physically located, altering the incident angles of the light rays on the object. Here, we present the ray tracing method to estimate

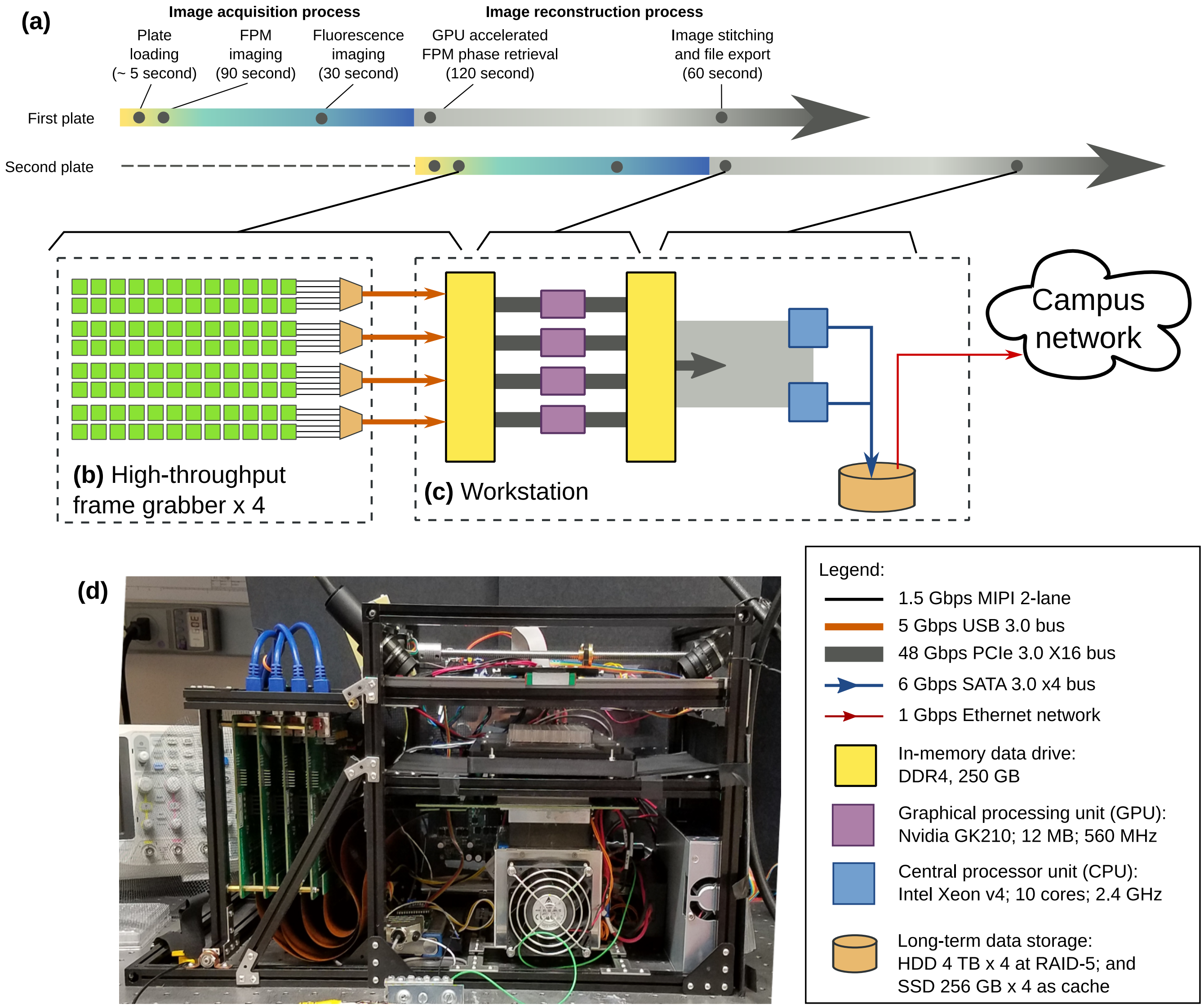


**Supplementary Figure S1.** **Parallel FPM acquisition and reconstruction process.**(a) Timeline of plate image acquisition and reconstruction processes two consecutive plates. Since the reconstruction process can be done offline, the second plate can be loaded and imaged while the workstation is reconstructing the images of the first plate. (b) and (c) Four (4) high-throughput frame grabbers streams raw images to the internal memory buffers of the workstation through the high speed links. (d) Front view of the 96 Eyes hardware.

the incident angles due to refraction.

First, we consider the case when the liquid interface is devoid of meniscus. Let us denote the vertical distance between the object and the light source by $h_a$, and the liquid medium (refractive index$= n$) height above the object by $h_b$. For a light ray from a single LED passing through the flat air-to-liquid interface [inset of Supp. Fig. S2], the angle of illumination on the sample $\theta$ is governed by

$$n \sin\theta = \frac{x_a - x_b - \delta}{\sqrt{(h_a - h_b)^2 + (x_a - x_b - \delta)^2}} \quad \text{Snell's law} \tag{S1}$$

$$\sin\theta = \frac{\delta}{\sqrt{h_b^2 + \delta^2}} \quad \text{Geometry} \tag{S2}$$

The close form solution of $\sin\theta$ exists, but it involves finding the root of a fourth order polynomial derived from the above equations. Instead, we numerically solve for $\delta$ with the following root-finding algorithm

$$\delta^0 = \frac{(x_a - x_b) h_b}{h_a} \tag{S3}$$

$$\delta^{k+1} = g(\delta^k)$$

$$\text{subject to } g(\delta) = \frac{1}{n}\left[\frac{(x_a - x_b - \delta)\sqrt{h_b^2 + \delta^2}}{\sqrt{(h_a - h_b)^2 + (x_a - x_b - \delta)^2}}\right], \tag{S4}$$

which guarantees to converge for $|x_a - x_b| < h_a - h_b$. The illumination angle can be now evaluated by substituting $\delta^K$ into Eq. S2 for a large number $K$.

Next, we analyze the changes to the optical light path in the presence of meniscus. The meniscus introduces a tilted air-to-liquid interface at an angle $\alpha(x_b)$, which is a function of the lateral position $x_b$ from the center of the well on the culture plate. Trying to incorporate this variable to the ray-tracing model will add unnecessary complexity to Eq. S1. Therefore, we linearize the meniscus effect by introducing a parallax shift $x_p$, with

$$x_p(x_b) = (h_a - h_b)[\tan(\alpha(x_b) + \theta) - \tan\theta] \approx c x_b, \tag{S5}$$

for some constant $c > 0$. The meniscus-compensated illumination angle $\theta$ is now approximated by modifying Eq. S4 with $x_a \mapsto x_a - x_p(x_b) \approx x_a - c x_b$.

### Speed improvement factor and the design criteria of the parallel illumination scheme

Without parallel illumination, only a single camera is active at any instance of image acquisition. Let $f$ be the effective frame rate of a single camera. For the 96-well plate, the total acquisition time required is equal to $f^{-1} \times$ number of wells $\times$ number of illumination $= 4704 f^{-1}$. Parallel illumination scheme instead utilizes a 2D lattice illumination pattern with a source-to-source separation of $m$ LEDs

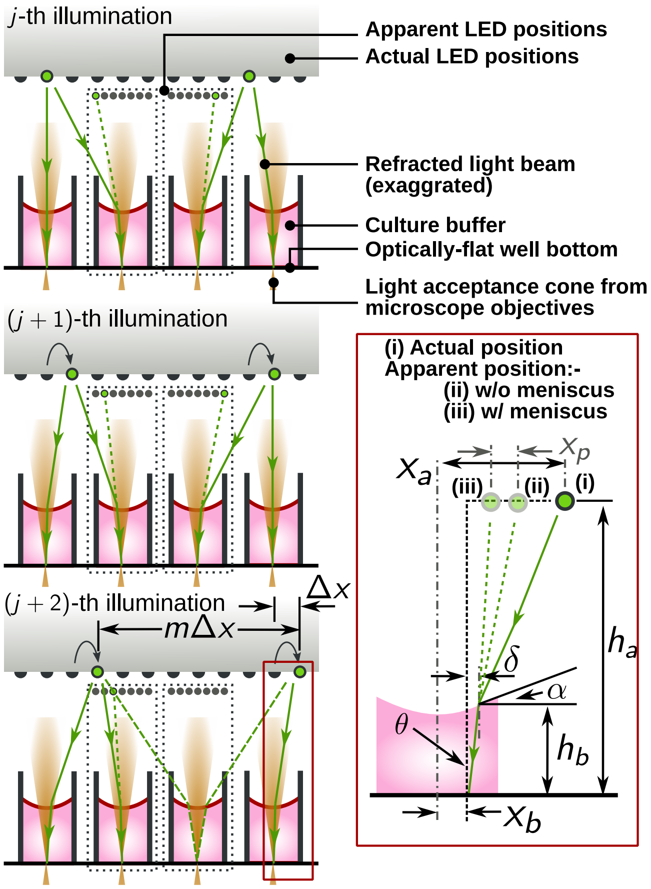


**Supplementary Figure S2. Detailed illustration of the parallel illumination scheme of 96 Eyes.** The source-to-source separation is chosen to maximize the effective acquisition rate, as well as avoiding interference. This is made possible by making sure that only one single LED is responsible for brightfield illumination for any camera and for any time instance of ptychographic image acquisition. Inset: definition of symbols for LED position calibration.

with a LED-to-LED separation of $\Delta x$. The total number of illumination is now reduced to $m^2$. Hence, the effective acquisition time is equal to $f^{-1} \times m^2 \times$ number of cameras $\times$ (number of frame grabber cards)$^{-1} = 24 m^2 f^{-1}$, for four frame grabber cards. The speed improvement is given as

$$\frac{4704 f^{-1}}{24 m^2 f^{-1}} = 196 m^{-2}. \tag{S6}$$

In the following paragraph, we will compute the lower limit of value $m$. For all time instances, we only allow one LED to fulfill the brightfield illumination condition with respect to the object (Fig. S2). Let us denote the vertical distance between the object and the light source by $h_a$, and the liquid medium (refractive index$= n$) height of $h_b$. The following conditions have to be fulfilled in addition to Eqs. (S1) and

(S2):

$$n \sin\theta \geq \text{NA} \tag{S7}$$
$$x_b = 0 \tag{S8}$$
$$x_a = m\Delta x/2 \quad \text{subject to } m = 2^M, \tag{S9}$$

for a given numerical aperture (NA) of the microscope objective, and some integer $M$. Here a power of two is preferred because it simplifies the electronic design of the LED matrix. For our system with $h_a = 33\,\text{mm}$, $h_b = 3\,\text{mm}$ and $\Delta x = 3\,\text{mm}$, we picked $m = 8$. This implies a conservative speed improvement of at least 3 times. Compared to mechanical scanning system which has a much lower effective frame rate $f$, the speed improvement can be up to 8 times compared to commercially available instruments.

### Modification to the Fourier ptychography phase retrieval algorithm

**Forward model for our imaging system** Let us denote a segment of the object to be reconstructed by $u \in \mathbb{C}^n$, a two-dimensional image with $n^{1/2} \times n^{1/2}$ pixels. We also denote the $j$-th illuminated low-resolution intensity image of the object by $I_j \in \mathbb{R}^m_+$, with $m^{1/2} \times m^{1/2}$ pixels ($u$ and $I_j$ are both written as a vector by a lexicographical order). It can be shown that $I_j = |\mathcal{F}^{\text{H}} \text{diag}(p) \mathbf{Q}_j \mathcal{F} u|^2$. The pupil function $p \in \mathbb{C}^m$ can be considered as the circular aperture at the back aperture plane of the imaging system. Binary matrix $\mathbf{Q}_j \in \mathbb{R}^{m \times n}$ depicts the downsampling of the object $u$ by cropping a region of $m$ pixels in Fourier space corresponding to the $j$-th position of the light source. What we measure is a stack of low-resolution intensity images $I_j = |\mathcal{F}^H \psi_i|^2 = |\mathcal{F}^H \text{diag}(p) \mathbf{Q}_j \mathcal{F} u|^2 \in \mathbb{R}^m_+$, $j = 1, 2, \ldots, k$, where the hyperscript $H$ denotes a Hermitian conjugate. The operation $\text{diag}(a) b$ represents the element-by-element multiplication[2] between two vectors $a, b$.

In reality, the measured sequence of low-resolution images are corrupted by (i) the ambient light level $I_b > 0$, (ii) angular dependency of LED intensity $w_j > 0$, (iii) background interference of suspended particulates in the liquid $\phi_{\text{dust}} \in \text{R}^n$; and (iv) dark current and readout noise of the sensor $n_j \in \text{R}^m$. Therefore, we modified the forward model to

$$I_j = w_j |\mathcal{F}^H \text{diag}(p) \mathbf{Q}_j \mathcal{F} \text{diag}(e^{i\phi_{\text{dust}}}) u|^2 + I_b + n_j, \tag{S10}$$

**Minimizer with partial spatial coherence constraint** Since the lens aberration is almost completely unknown, one has to solve a blind ptychographic phase retrieval problem with an amplitude constraint[2]

$$\min_{\{w_j\}, p, u} \sum_{j=1}^{N} f_j(w_j, p, u)$$
$$\Leftrightarrow \min_{\{w_j\}, p, u} \sum_{j=1}^{N} \left\| |\mathcal{F}^H \text{diag}(p \sqrt{w_j}) \mathbf{Q}_j \mathcal{F} u| - \sqrt{I_j - I_b} \right\|_2^2, \tag{S11}$$

**Algorithm 1** Pseudo-code of the phase retrieval algorithm for 96 Eyes system

1. **Inputs:** segments of low resolution images $I_j$ and ambient light level $I_b$ of the corresponding camera.
2. Initialize local pupil functions $p_\ell$ for all $L$ segments of the object.
3. Estimate the global pupil function $\sum p/L := (1/L)\sum_{\ell=1}^{L} p_\ell$.
4. **FPM-EPRY algorithm:** Run the phase retrieval algorithm for the $\ell$-th segment with pupil function recovery.
   (a) **Initialize** $u^0 := \sqrt{I_0 - I_b}$, $p^0 := \sum p/L$ and $w_j^0 := 1$ for $j \in [1, N]$.
   (b) For the $k$-th iteration,
      i. Evaluate $j = \text{mod}(k, N) + 1$.
      ii. **Object update:** solve $u^{k+1} = \arg\min_u f_j(w_j^k, p^k, u)$.
      iii. **Weighting update:** when $k \leq 3N$, solve $w_j^{k+1} = \arg\min_w f_j(w, p^k, u^{k+1})$. Otherwise, $w_j^{k+1} = w_j^k$.
      iv. **Pupil update:** when $k > 3N$, solve $p^{k+1} = \arg\min_p f_j(w_j^{k+1}, p, u^{k+1})$. Otherwise, $p^{k+1} = p^k$.
   (c) Repeat step (4b) until $k = K$.
   (d) Update the object estimate $u_\ell := u^K$ and local pupil function estimate $p_\ell := p^K$.
5. Repeat steps (3)–(4) one more time.
6. **Separate $u$ from $e^{i\phi_{\text{dust}}}$** by digitally high-passing the phase component of $u_\ell$ with an inverted Gaussian blur kernel. The amplitude component is preserved as $|u_\ell^{\text{cell}}| := |u_\ell|$.
7. **Stitch** the recovered image segments $u_\ell$ for all $\ell \leq L$.
8. **Outputs:** amplitude and phase component of the stitched image $u$, the global pupil function $\bar{p}$ and the local aberrations $\{p_\ell\}$.

Because of the limited number of low-resolution images (=21) in the measurement, the estimated pupil function $p^{\mathrm{est}}$ cannot be efficiently separated from the estimated object $\mathcal{F}u^{\mathrm{est}}$in the Fourier domain. This shortcoming is compounded by the fact that the target biological specimen is a weak phase object, where most of the information in the Fourier domain is concentrated in that of the un-scattered transmitted light. To suppress the crosstalk between the two, we utilize a finite number ($L > 0$) of overlapping segments of the object $u_\ell$ and the corresponding local pupil $p_\ell$ to enforce the partial spatial coherence constraint. That is, the above minimizer is further subject to

$$\sum_{\ell=1}^{L} \| p_\ell - \sum p/L \|_2^2 \leq \epsilon^{\mathrm{tol}}, \tag{S12}$$

for a "global" average pupil function $\sum p/L = (1/L)\sum_{\ell=1}^{L} p_\ell$ and tolerance value $\epsilon^{\mathrm{tol}} > 0$.

**Background estimation** To recover the average level of the ambient light level $I_b$, we capture the images when all light sources are switched off. The value of $I_b$ for a particular CMOS sensor is then set to be the pixel average of the captured dark image.

**Separation of the non-uniform illumination profile of LEDs and the pupil function** From Eq. (S11), it is known that the pupil function $p$ cannot be efficiently separated from the factor $w_j$. Therefore, the factor $w_j$ is optimized only for the first three iterations[3], while the recovery of $p$ is postponed until the fourth iteration.

**Separation of cells and background interference** The out-of-focus suspended particulates show up as blurred shadows in the sequence of low-resolution images [Supp. Fig. S3]. We utilize this property to estimate $\phi_{\mathrm{dust}}$ by applying a Gaussian blur of the recovered object phase. The morphological information of the cells can be extracted from the phase difference between the recovered field $u^{\mathrm{est}}$ and $e^{i\phi_{\mathrm{dust}}}$.

It is noted that there are existing algorithms that specializes in separation of the object from out-of-focus noise[4].

**Choice of adaptive step size for pupil recovery** While the object update in Step 4(b)ii of Algorithm 1 is solved by the time-honored Gaussian-Newton algorithm[5], the pupil update in Step 4(b)iv of Algorithm 1 is instead solved by the gradient descent method[6], with

$$\begin{aligned} p^{k+1} &= \arg\min_p f_j(w_j^k, p, u^k) \\ &= p^k + \gamma \operatorname{diag}(\bar{s}^k) \times \\ &\quad \left[ \mathcal{F}\operatorname{diag}\left( \frac{\sqrt{I_j}}{|\mathcal{F}^H g_j(w^k, p^k, s^k)|} \right) \mathcal{F}^H g_j(w^k, p^k, s^k) - \right. \\ &\quad \left. g_j(w^k, p^k, s^k) \right], \end{aligned} \tag{S13}$$

where $s^k = \mathcal{F}u^k$ and $g_j(w^k, p^k, s^k) = \operatorname{diag}(p^k \sqrt{w_j^k})\mathbf{Q}_j s^k$ for a step size of $\gamma \in \mathbb{R}_+^m$. Because of the choice of parallel illumination in our 96 Eyes system, all of the captured data are brightfield images. For a weak phase object, most of the incoming light rays remains un-scattered, that result in a strong peak in the Fourier domain. If the step size $\gamma$ is a constant, the recovered pupil will be corrupted with a constellation like artifact [Fig. S4(a)]. Therefore, we heuristically adjust the step size with

$$\gamma = \left[ \operatorname{diag}\left( (1-\beta)|s^k| + \beta \|s^k\|_\infty \right) \right]^{-1}, \tag{S14}$$

where $\|s\|_\infty$ denotes the maximum amplitude of the complex-valued signal $s$. Effectively, the step size $\gamma$ normalizes the value of $\operatorname{diag}(\bar{s}^k)$. The non-dimensional number $\beta \in [0, 1]$ adjusts the relative strength of normalization of signal $\bar{s}^k$. When $\beta = 0$, the Fourier domain of the object $s^k = \mathcal{F}u^k$ will be completely normalized. In the main text, the value is set to be $\beta = 10^{-6}$, around twice the order-of-magnitude of an 8-bit image. This helps smooth the pupil function [Fig. S4(b)] as well as reduce the reconstruction residual [Fig. S4(c)], defined

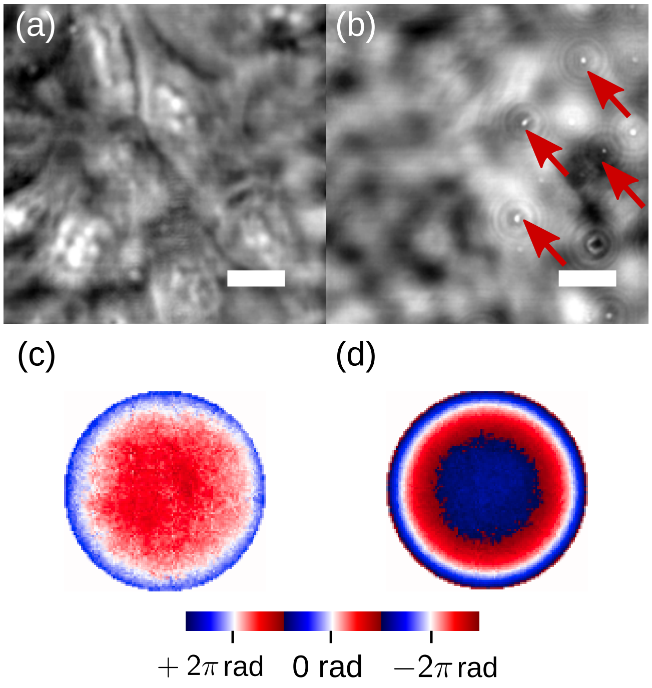


**Supplementary Figure S3. Particulates outside of the focal plane introduce background interference.** (a) Phase component of the recovered complex wavefront, showing the the U2OS cell line almost buried in the phase fluctuation; (b) phase image of dust particles on the underside of the well plate, reconstructed by digital refocusing of the recovered complex wavefront.
(c) Recovered phase component of system aberration; and (d) pupil function used for digital refocusing.

as

$$\epsilon^k = \frac{\sum_{j=1}^{N} \left\| |\mathcal{F}^H g_j(w^k, p^k, s^k)| - \sqrt{I_j - I_b} \right\|_2^2}{\sum_{j=1}^{N} \| \sqrt{I_j - I_b} \|_2^2}. \quad \text{(S15)}$$

### Improving the dynamic range of fluorescence images with two-stage digital averaging

Because of the limited photo-sensitivity and bit depth of our choice of consumer-grade CMOS sensor, we adopted the digital averaging approach to enhance the signal-to-noise ratio of the sensor. The digital averaging technique is also known as *dithering* in audio digitization community[7–9], where the band-limited signal of interest is mixed with an artificial out-of-band noise on the input side of the analog-to-digital conversion circuit to reduce the quantization error. Our technique is also very similar to *halftoning* of digital images[10], where an artificial pepper noise is added to simulate grayscale images out of a black-and-white display device. In contrast,

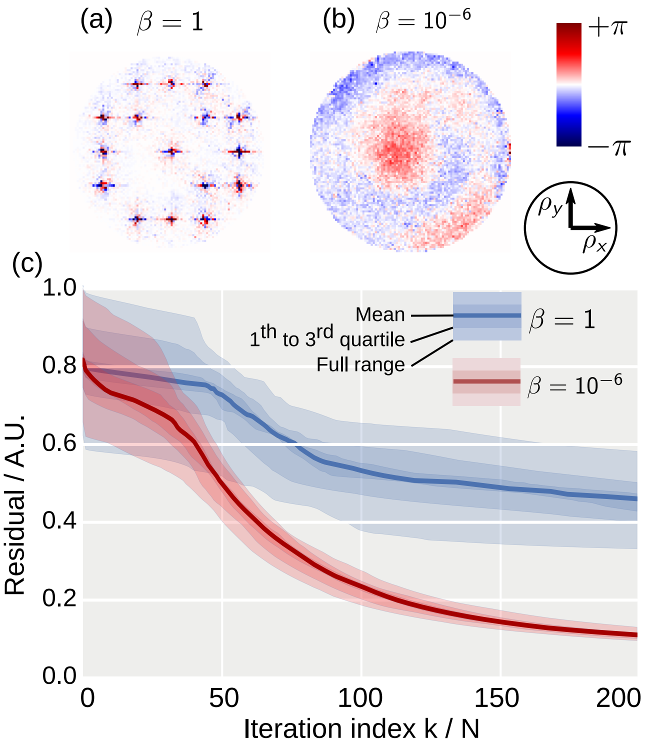


**Supplementary Figure S4. Adaptive step size improves pupil function recovery.** (a) Recovered phase component of pupil function with a constant step size, i.e. at $\beta = 1$, compared to (b) at $\beta = 10^{-6}$. Symbols $(\rho_x, \rho_y)$ are the local coordinates of the pupil function. (c) Comparison of reconstruction residuals by applying phase retrieval to all segments ($L = 80$) of the cell sample captured from one camera. With our method, the residual reduces by around one-third (after $k/N = 200$ iterations) with a much smaller spread, demonstrating a more robust object and pupil co-recovery.

the noise source for our CMOS sensors in the 96 Eyes system cannot be precisely controlled. Notably, similar digital averaging approaches has been proposed before for radiometry studies[11]. However, the underlying principle is poorly understood. Here, we provide a theoretical framework to offer to explain the dynamic range improvement of our fluorescence images with digital averaging.

**Forward model** For a fluorophore concentration $c(x,y)$ illuminated by an uniform intensity $I_0$, the imaging system in the fluorescence channel is empirically modeled as

$$\begin{aligned} I(x,y,t) &= \lfloor g_{\text{amp}} \eta c(x,y) I_0 + g_{\text{amp}} n_{\text{dark}}(x,y,t) + n_{\text{amp}}(t) \rfloor \\ &= g_{\text{amp}} \eta c(x,y) I_0 + n_{\text{amp}}(t) + \\ &\quad g_{\text{amp}} n_{\text{dark}}(x,y,t) + \epsilon(x,y,t), \end{aligned} \quad \text{(S16)}$$

where the non-dimensional factor $\eta$ is a product of (i) quantum efficiency of the fluorophore, (ii) photon collection efficiency of the microscope objectives, and (iii) quantum efficiency of the photosensing circuit in the CMOS sensor. The amplifier with gain $g_{\text{amp}} > 0$ naturally comes with an additive power-line noise $n_{\text{amp}}(t)$. The round-off operator $\lfloor \cdot \rfloor$ denotes the quantization process, which in turn can be modeled as an additive quantization error $\epsilon(x,y,t) \in [-0.5, +0.5)$. Here, the photon noise is assumed to be negligible compared to dark current noise.

In rolling shutter mode, rows of pixels are read out at a traversal rate of $v$, so the amplifier noise is mapped to the vertical axis of the $j$-th image $I_j(x,y)$.

$$\begin{aligned} I_j(x,y) &= g_{\text{amp}} \eta c(x,y) I_0 + n_{\text{amp}}(y,j) + \\ &\quad g_{\text{amp}} n_{\text{dark}}(x,y,j) + \epsilon_j(x,y), \end{aligned} \quad \text{(S17)}$$

where $n_{\text{dark}}(x,y,j) = n_{\text{dark}}(x,y,t)|_{t=t_j+y/v}$ and $n_{\text{amp}}(y,j) = n_{\text{amp}}(t = y/v + jH/v)$ for $H$ rows of pixels in the CMOS sensor.

**Suppressing both dark current noise and quantization error with digital averaging** Consumer-grade CMOS sensors, designed for daylight applications, have a much higher quantization error than the dark current noise. For a amplifier gain value $g_{amp}$ at unity, the dark current noise component is typically always rounded-off to zero. In other words, direct digital averaging of multiple frames $I_j$ at unity gain usually do not result in reduction of quantization error. However, typical biological specimen is known to have a low fluorophore concentration. Concerns about photobleaching also limit the illumination intensity $I_0$. Therefore, the amplifier gain $g_{\text{amp}}$ must be boosted sufficiently to utilize the full quantization range of the CMOS sensor.

The dark current noise is known to possess a Gaussian distribution[12], i.e. $n_{\text{dark}}(t) \sim \mathcal{N}(0, \sigma^2_{\text{dark}})$ (symbols $x,y$ are omitted for clarity). This also applies to the power-line noise, where $n_{amp}(t) \sim \mathcal{N}(0, \sigma^2_{\text{amp}})$. For a sufficient gain with $g\sigma_{\text{dark}} \gg 0.5$, the probability density function of $I_j$ is given as

$$P(I_j = a) = \begin{cases} \frac{1}{A}\int_{a-0.5}^{a+0.5} \exp\left(\frac{-(I-g_{\mathrm{amp}}\eta c I_0)^2}{(\sigma^2_{\mathrm{amp}}+g^2_{\mathrm{amp}}\sigma^2_{\mathrm{dark}})\sqrt{2}}\right) dI & \text{if } a \text{ is an integer,} \\ 0 & \text{otherwise.} \end{cases} \tag{S18}$$

The scaling factor $A$ is defined such that $\int_{-\infty}^{\infty} P(I_j = a)da = 1$. By averaging a sufficient number of frames, i.e.

$$I_1(x,y) := \frac{1}{N}\sum_{j=1}^{N} I(x,y,t_j), \tag{S19}$$

both the dark noise and the quantization error can be reduced. For instance, it can be shown that $\lim_{N\to\infty} I_1(x,y) = \int_{-\infty}^{+\infty} a\,P(I_j = a)\,da = g_{\mathrm{amp}}\eta c(x,y) I_0$, which is independent of both noise terms.

**Suppressing the power-line noise** For our digital-averaged fluorescence signal captured by the 96 Eyes system, we can still observe the presence of row-wise intensity fluctuation [Supp. Fig. S5(b)] originated from the power-line noise above the quantization level, modeled as $I_1(x,y) \approx g_{\mathrm{amp}}\eta c(x,y) I_0 + n_{\mathrm{amp}}(y)$. This is caused by the power-line noise in the 96-in-1 camera board.

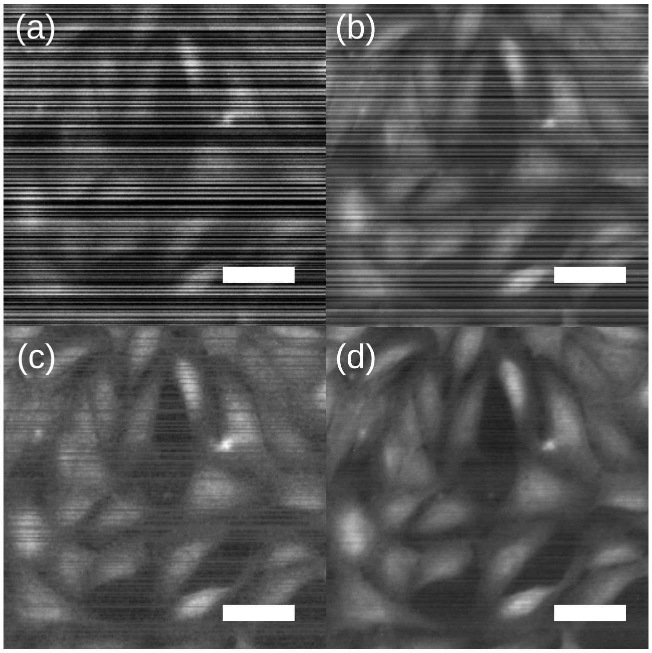


**Supplementary Figure S5. Improving the dynamic range of fluorescence images with digital averaging.** (b) Single frame at gain $g_{\mathrm{amp}} = 8$; (c) averaging 10 frames at gain $g_{\mathrm{amp}} = 8$; Suppressing the band-like pattern noise for (c) a single frame, and (d) the digital average of 10 frames. All images are contrast-stretched to highlight the background noise and artifacts. Scale bar: 20 µm.

The size constraint of the 96-well culture plate limits the available real estate on the printed-circuit board for electronic filters, especially the decoupling capacitors. To further suppress such row-wise fluctuations, we apply the same digital averaging technique to isolate it from the fluorescence signal $c(x,y)$. Here, we assume that the fluorophore concentration possesses a Gaussian distribution $c(x,y) \sim \mathcal{N}(\mu_c, \sigma_c^2)$ for $\mu_c > 0$, $\sigma_c \ll \mu_c$. By taking a row-wise average of pixels of $I_1(x,y)$, we have

$$I_2(y) := \frac{1}{W}\sum_{i=1}^{W} I_1(x_i,y) \approx g_{\mathrm{amp}}\eta\mu_c I_0 + n_{\mathrm{amp}}(y), \tag{S20}$$

for $W$ pixels along individual rows of the image. Since we only care about the morphology of the biological cells stained with the fluorophore, the average fluorophore concentration $\mu_c$ can be eliminated as well. Hence, the recovered fluorescence image is given as

$$c^{\mathrm{est}}(x,y) := c(x,y) - \mu_c \approx \frac{I_1(x,y) - I_2(y)}{g_{\mathrm{amp}}\eta I_0}. \tag{S21}$$

In practice, the signal $c(x,y)$ does not fit well with the Gaussian process assumption. The row-wise averaging operation Eq. S20 is replaced with row-wise median operation to reduce sensitivity to extreme values in $c(x,y)$.

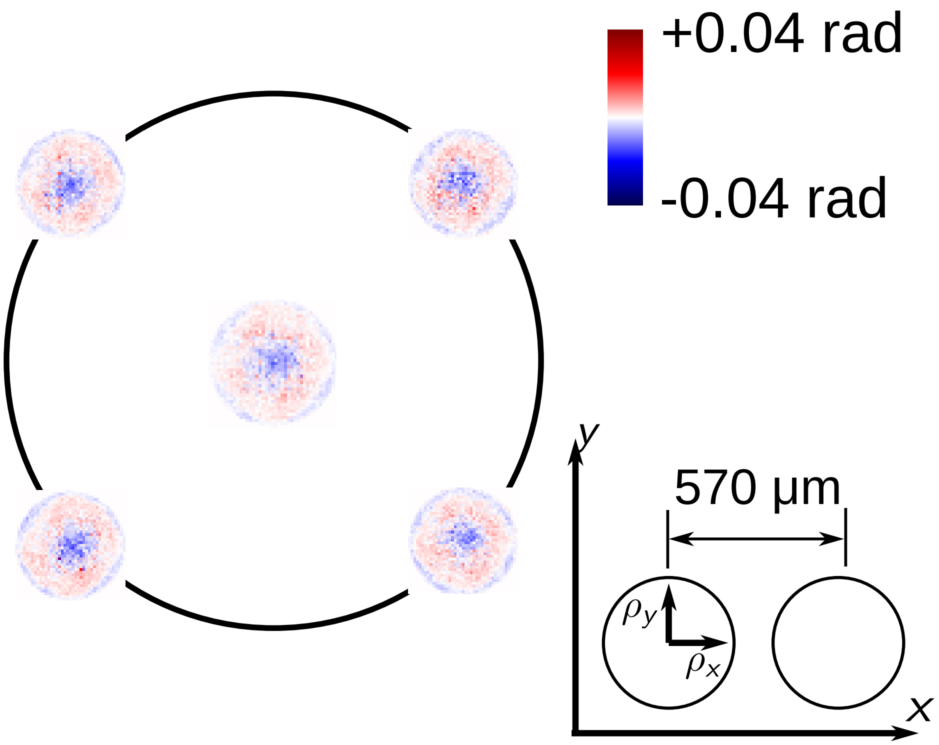


**Supplementary Figure S6.** **Surface flatness of the polystyrene samples, analyzed by the scientific-grade FPM.**

**12.** Nakamura, J. *Image Sensors and Signal Processing for Digital Still Cameras* (CRC Press, 2005).

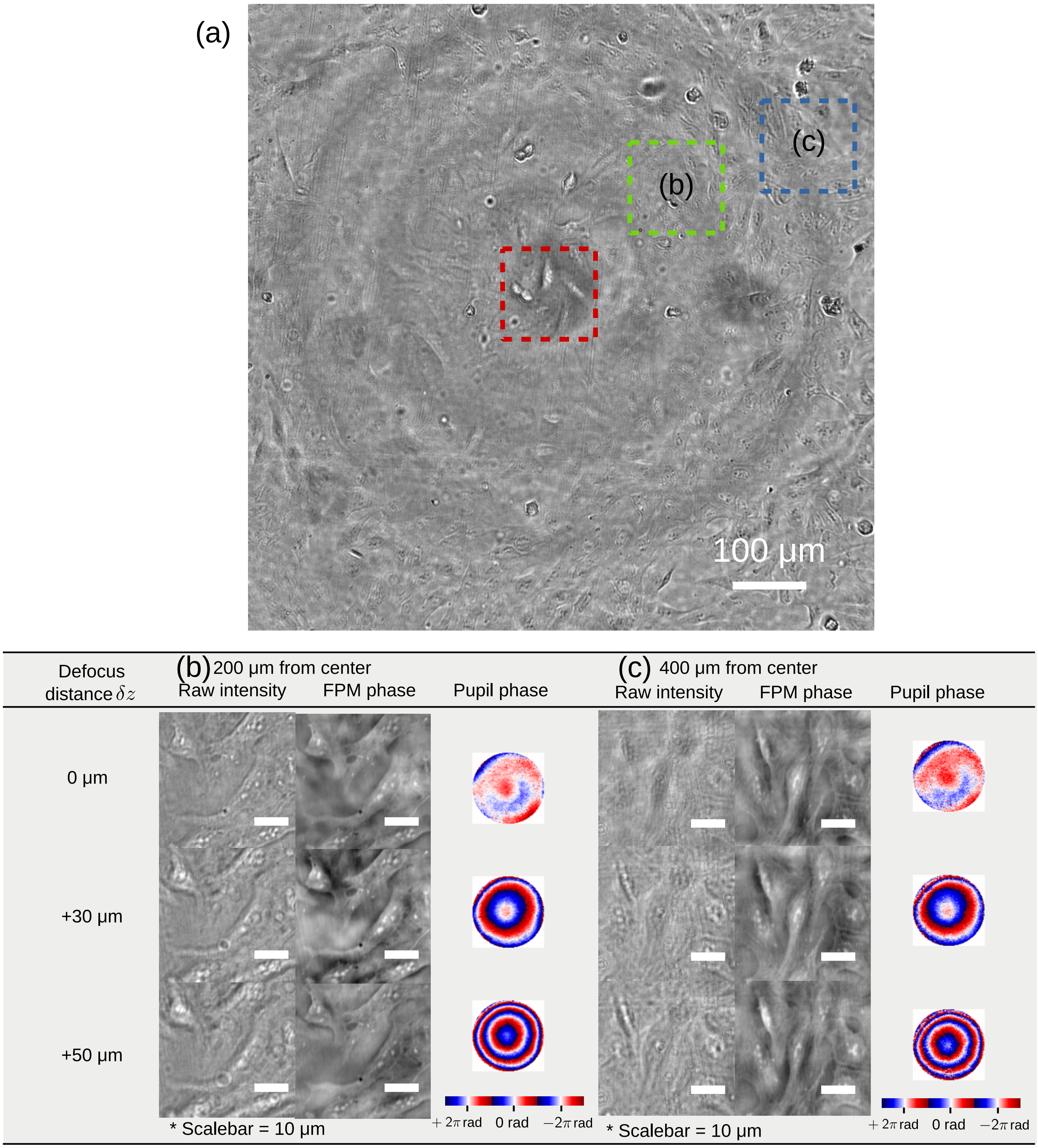


**Supplementary Figure S7. Computationally refocused phase images at off-axis locations.** (a) Raw intensity image of the entire field-of-view of the U2OS cell line. Also shown are the FPM Phase reconstruction (b) halfway from the edge of the field-of-view; and (c) close to the edge of the field-of-view.

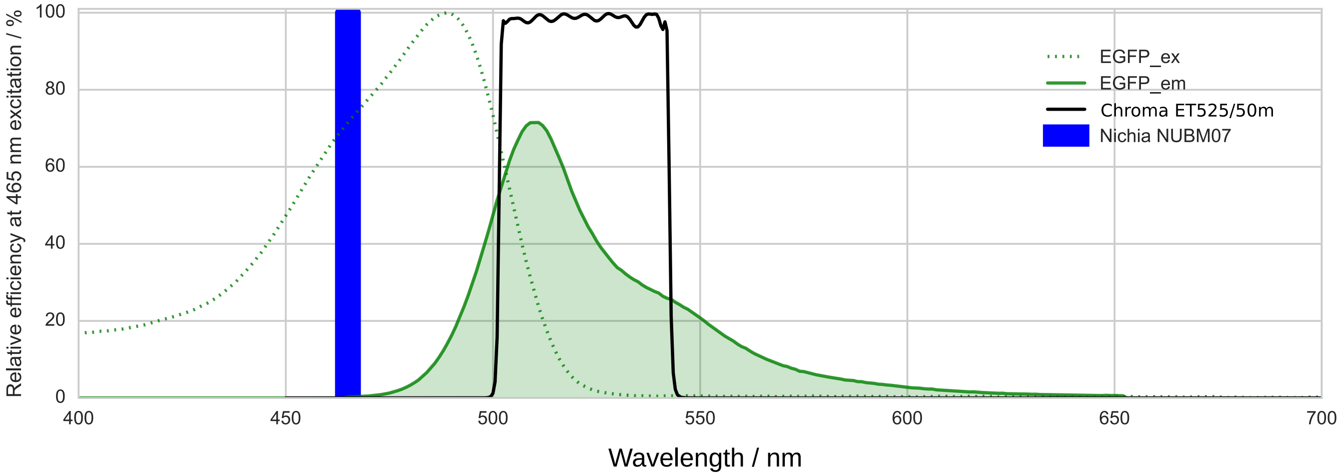


**Supplementary Figure S8. Spectra of laser, fluorophore (eGFP) and filter set for fluorescence microscopy.** The multimode diode laser (Nichia NUBM07) is filtered with a laser clean up filter of a 5 nm bandwidth (Semrock FF01-465/5).
